# Functional and Structural Brain Age Gaps as Complementary Predictive Markers for Cognitive Performance

**DOI:** 10.64898/2026.10.20.760445

**Authors:** Annachiara Crocetta, Marta Morandini, Nora Bittner, Svenja Caspers, Franco Cauda, Andrea Brovelli, Matthieu Gilson

## Abstract

Cognitive decline is characterized by substantial inter-individual variability across the lifespan, even among clinically healthy adults. However, it is unclear how structural and functional brain organization jointly relate to differences in cognitive performance within the typical populations. We considered a multimodal brain age framework by predicting for each subject its age from either structural MRI or resting-state functional MRI, after training on the 1000BRAINS cohort (901 participants, 18–85 years). Following, a structural and a functional brain age gap markers were defined as the deviations with respect to the real chronological age. We then used the markers to characterize the interplay between structure and function in shaping cognitive performances. Our findings reveal that the functional marker is as important to predict global cognitive performance as the structural marker. The most accurate prediction was found for global cognition, executive functions and episodic memory, followed by working memory, attention and language. In particular, the structural marker was contributing to all domains, except for working memory, whereas the functional marker had the highest contribution. Overall, structural and functional brain age models together provide a meaningful subject-level summary of brain information for predicting cognition, supporting a multidimensional view of typical aging.

## Introduction

The rise in life expectancy has reshaped the demographic structure of modern societies, making age-related disorders a priority for public health^1^. Cognitive decline toward pathological conditions, such as dementia or other neurodegenerative diseases, is particularly relevant, because of personal, societal and economic consequences^2–4^. However, the characterization of cognitive aging remains an open issue, due to high inter-individual variability in cognitive profiles among individuals of similar chronological age^5,6^, and various sociodemographic factors, such as education, socioeconomic status and lifestyle factors^7,8^. Explaining this heterogeneity is a central challenge for human lifespan neuroscience as age-related cognitive evolution is likely affected by the interplay of multiple structural and functional processes. In particular, the mechanisms governing their unfolding in time in cognitive decline remain poorly characterized^9^.

Within the neuroimaging field, magnetic resonance imaging (MRI) provides a non-invasive in vivo approach for characterizing brain mechanisms underlying individual differences in cognitive aging. Recent studies suggest that brain trajectories may be better understood by jointly considering brain structural and functional phenotypes. First, brain trajectories have been shown to be organized along a macroscale cortical gradient, the so-called sensorimotor–association (S-A) axis^10,11^, which ranks cortical areas from primary, sensory and motor areas to higher-order associative cortices, and reflects differences in cytoarchitecture, myelination, gene activity, and functional connectivity^12,13^. In fact, the brain develops along the S-A axis^11^, with association areas maturing last and declining first^14^, and regional patterns of accelerated brain aging also align with it^15,16^. Second, distinct structural and functional brain trajectories may be associated with preserved or altered cognitive performance^17–21^. Note that while structural brain measures provide an important index of age-related cognitive changes and neurodegenerative diseases^22–24^, brain structure alone is not the sole determinant of cognitive abilities^25,26^. Cognitive performance during aging reflects the complex interplay between structural and functional organization with network-level reconfiguration^27–29^, rather than either factor alone^30,31^. Accordingly, recent works have highlighted the advantage of multimodal approaches for capturing how anatomical and functional variation jointly shape cognitive heterogeneity^20,32–34^, helping identify early signatures of vulnerability, as well as trajectories of preserved functioning and resilience^32,35^.

Lately, the Brain Age Gap (BAG) framework has emerged as a powerful approach for quantifying individual deviations from expected population average^36–38^. Calculated as the difference between the age predicted from brain imaging techniques and the chronological age, the BAG provides a subject-specific marker of *biological* brain age, quantifying whether the brain appears older or younger than expected. BAG has been widely applied to neurodegenerative disease, such as Alzheimer’s disease and frontotemporal dementia^39,40^, and recent meta-analytic evidence further supports its relevance across a broader range of neurological and psychiatric conditions^41^. BAG is also informative in cognitively healthy populations^40,42^. In fact, the BAG has been associated with a broad range of health, demographic, lifestyle-related risk factors such as smoking^43^, mortality^44^, social disadvantage^45,46^, and protective factors such as physical activity^47^, and childbirths^48^.

While anatomical models achieve high accuracy in predicting chronological age from structural imaging features^49,50^, functional models may capture inter-individual variation relevant for clinical and behavioural phenotypes, such as anxiety, depression, subjective cognitive complaints and mild cognitive impairment^20,51^. Since functional connectivity alterations often precede detectable structural decline^52^, the integration of functional data with anatomical features could enhance the detection of cognitive impairment^34,53^. More recently, multimodal BAG approaches have started to move beyond structural MRI by incorporating additional MRI-derived measures^54^. This has paved the way for an in-depth investigation of resting-state functional organization that remains less well characterized. However, it remains unclear how structural and functional brain-age patterns jointly relate to this variability in typical aging.

In the current study, we investigate the extent to which inter-individual variability in typical ageing is reflected by structural and functional MRI patterns. To do so, we developed a multimodal BAG pipeline combining structural MRI and resting-state functional MRI to predict cognitive scores from the population-based 1000BRAINS cohort^55^. Our primary objective is to investigate whether structural and functional patterns of brain organization reflect cognitive heterogeneity across the lifespan. We further investigate whether high-dimensional structural and functional MRI information can be summarized into a lower dimensional subject-specific BAG marker that remains relevant for cognitive performance, beyond their role as measures of chronological age prediction.

Overall, the current work aims to refine the characterization of typical aging as a multidimensional continuum and to clarify how multimodal brain phenotypes may help identify vulnerability and preserved functioning before overt pathology emerges.

## Results

### Cognitive profiles reveal a hierarchical organization of structural and functional brain differences

We first investigated whether cognitive profiles were associated with distinct structural and functional brain signatures, by analyzing which phenotype is most sensitive to inter-individual differences in global cognition. Additionally, we examined whether this sensitivity is spatially organized along the S-A axis. By considering both modalities on a common cortical coordinate system, we can directly compare their cognition-related brain maps. Concordant alignment between modalities with the cortical axis would suggest largely overlapping information. On the contrary, different alignment might indicate a complementary effect, supporting structure and function as distinct brain-age markers in subsequent analyses.

In order to capture cognitive performance, we first analyzed the set of cognitive tests of the 1000BRAINS dataset, as illustrated in Fig. 1a. Cognitive assessments span a wide range of cognitive functions including executive functions, episodic and working memory, attention, and language abilities^17,55^. To extract the global cognitive score (GCS), we considered the first principal component across all tests, explaining 40% of the total variance (see Supplementary Section 2 for details on principal components analysis).

**Fig. 1:**
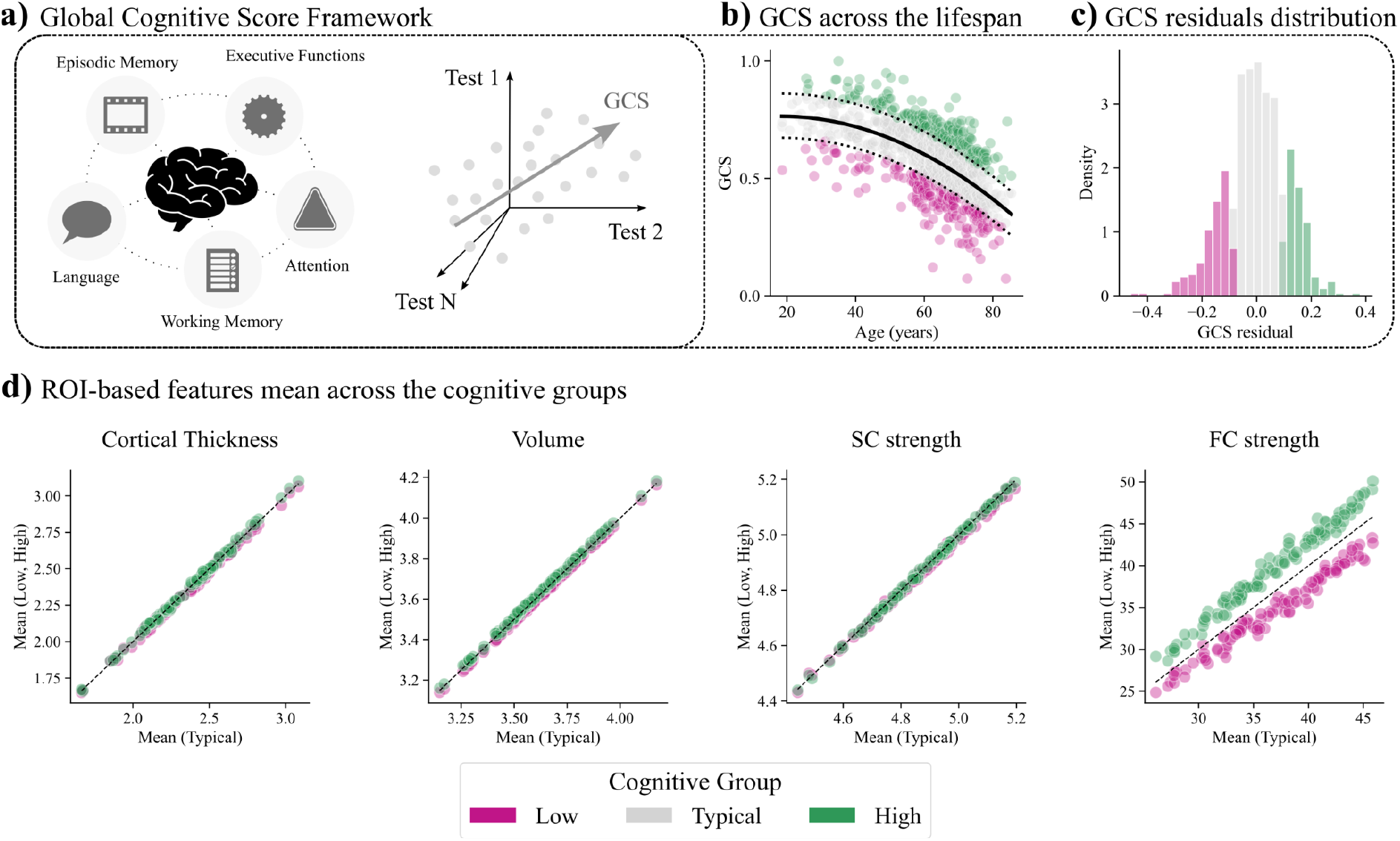
Relation of cognitive profiles to regional structural and functional brain phenotypes. **a)** Global Cognitive Score (GCS) Framework. Left: cognitive tests spanning different cognitive domains: executive functions, attention, language, episodic memory and working memory. Right: schematic representation of the GCS dimensionality reduction in the cognitive tests space. Dimensionality reduction is performed through principal component analysis (PCA). First PC explains 40% of the total variance. **b)** GCS across the lifespan stratified by the 20th and 80th percentiles, defining the Low, Typical and High cognitive performance profile groups (in pink, grey and green, respectively, see legend at the bottom of the figure). **c)** GCS residuals distribution from the predicted mean. **d)** ROI-based features mean in high (green) and low (pink) cognitive groups with respect to the typical (dashed line) cognitive profile: cortical thickness (p = 1, rb = -0.05), volume (p = 1, rb = -0.07), SC strength (p = 1, rb = -0.02), and FC strength (p < 0.001, rb = -0.51). The ROIs correspond to the Schaefer parcellation with 100 cortical regions.

To define cognitive profiles, we stratified participants using the 20th and 80th percentiles of the GCS quadratic regression^56^, thus resulting in three cognitive profiles: lower (below the 20th percentile, 181 participants), typical (between the 20th and 80th percentiles, 539 participants), and higher (above the 80th percentile, 181 participants) cognitive performance (Fig. 1b,c). This stratification was used to characterize group-level brain differences between the extremes of cognitive variability.

#### Relation between cognitive profiles and regional structural and functional brain phenotypes

We first studied whether the three cognitive profiles differ at the whole-brain level by considering the group-average values per region of interest (ROI) for the following brain phenotypes: cortical thickness, volume, structural connectivity (SC) strength and functional connectivity (FC) strength. All brain phenotypes are parcelled into the Schaefer 100 ROIs atlas^57,58^. See Methods for details. FC strength revealed the clearest mean difference between extreme cognitive profiles (p < 0.001, rb = -0.51), whereas all structural measures showed weaker group-level separation (Fig 1d). Differences were found to be consistent across age bins, and sex stratification (see Supplementary Section 3). Results suggest that functional organization may be sensitive to cognitive-profile differences. At the same time, given the weak relationship between structural phenotypes and cognitive profiles, we must go beyond group-level differences, taking into consideration the variability of global cognition.

#### The hierarchical organization of global cognition

We next considered the cognitive continuum, by examining the Spearman’s correlation (r) between regional features and the GCS, so as to better characterize the association between global cognition and brain features. Positive correlation indicates that higher cognition is associated with higher phenotype values (e.g. higher thickness), while negative correlation indicates higher global cognition when phenotype values are weaker (e.g. lower thickness).

Cortical thickness did not show a significant association with the S-A axis (r = 0.05, p_spin_ = 0.387), while volume (r = 0.41, p_spin_ < 0.01), SC strength (r = 0.49, p_spin_ < 0.001) and FC strength (r = -0.69, p_spin_ < 0.001) reveal distinct hierarchical organizations related to cognitive profiles (Fig 2a,b). In detail, volume and SC strength differences are stronger toward associative areas, usually considered associative hubs responsible for higher-level integration, in line with the hypothesis that structural vulnerability related to cognition might be particularly expressed in integrative systems^14^. Major differences arise from the ventral-attention (VAN), default mode (DMN), and fronto-parietal (FPN) networks, as shown in Fig. 2a,c. This may reflect the “last-in, first-out” (or retrogenesis) process^59^, a hypothesis in which the last areas to develop are the ones that degrade earlier and faster. On the contrary, all regions seem to drive the differences in cognitive profiles in FC strength, unveiling a strong negative association with the S-A axis. In fact, major differences are found to arise from regions belonging to the unimodals and attentional networks: somatomotor (SMN), visual (VN) dorsal attention (DAN) and ventral attention (VAN) networks, as shown in Fig. 2a,c. Results suggest a potential complementary mechanism or distinct biological pathways for structural decay versus functional reorganization, yet to be proven by taking into account age-related changes. Note that compared to current univariate analysis, multivariate regression of GCS from the different modalities did not show any association with the S-A axis (see Supplementary Section 5.3).

**Fig. 2:**
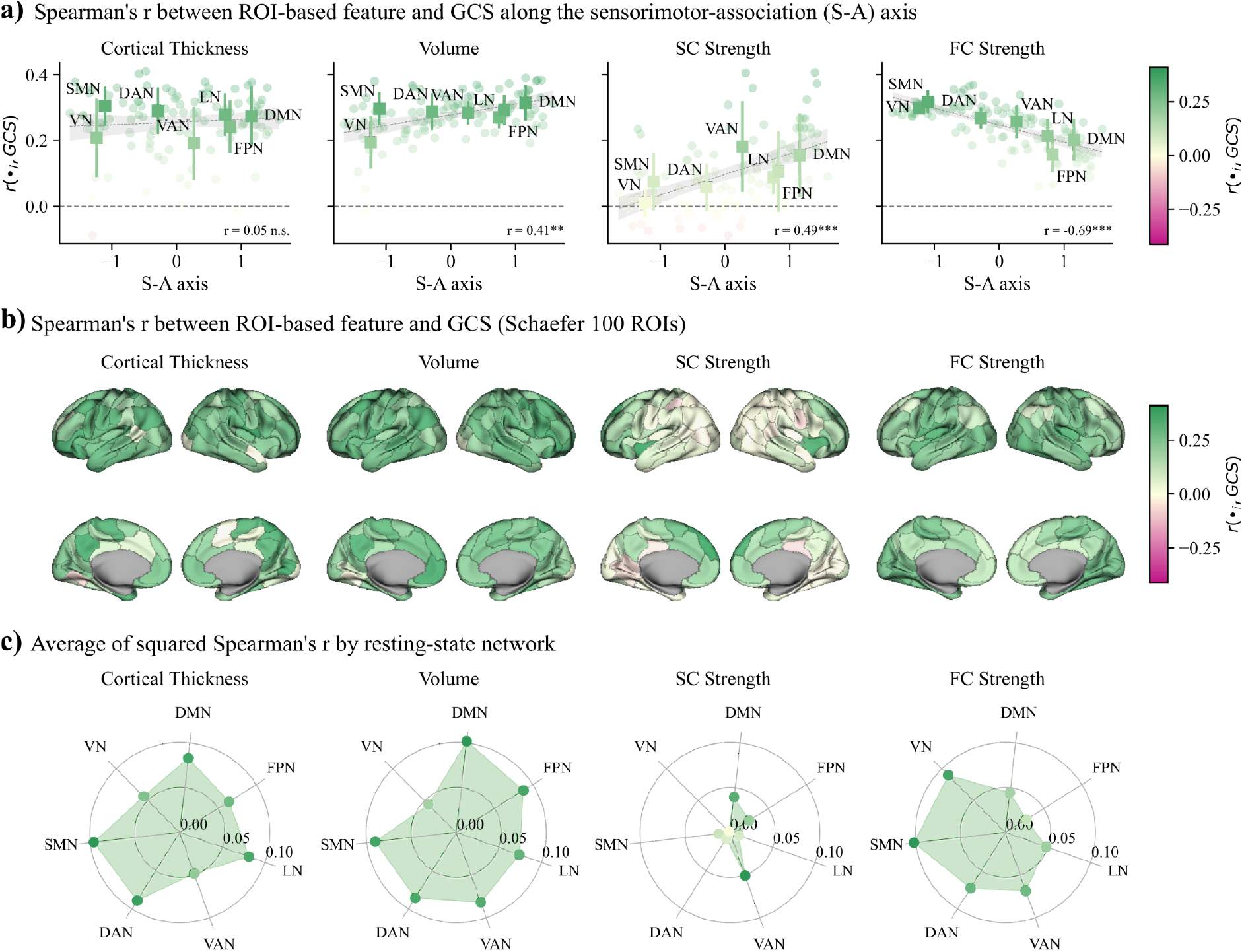
The hierarchical organization of global cognition. **a)** ROI-based Spearman’s correlation with GCS along the sensorimotor-association (S-A) axis. The point plot represents the average correlation values (square markers) per resting state network, with their respective standard deviation (error bar). S-A axis shows no significant association with cortical thickness (r = 0.05, p_spin_ = 0.387), while it is significantly associated with volume (r = 0.41, p_spin_ < 0.01), SC strength (r = 0.49, p_spin_ < 0.001), and FC strength (r = -0.69, p_spin_ < 0.001). **b)** ROI-based Spearman’s correlation with GCS on the 100 ROIs Schaefer atlas. Positive values (greener) indicate positive correlation with GCS. Negative values (pinker) indicate negative association between the ROI-feature and GCS. **c)** Average of squared Spearman’s r per resting-state network. Cortical thickness and volume have high association with GCS across most RSNs, while VAN, DMN and FPN seem to drive the SC strength association with GCS. Sensory and attentional RSN show greater association between FC strength and GCS. n.s.: non significant, *: p < 0.05, **: p < 0.01, ***: p < 0.001.

### Structural and functional brain age gaps show distinct cortical signatures

While regional brain associations with global cognition provide a lens into typical cognitive decline, understanding how brain structure and function interact at the subject-specific level remains challenging. In order to characterize the relation between age-dependent variability in brain structural and functional measure and cognitive performance, we next adopted a Brain Age Gap (BAG) framework. We defined for each subject two BAG markers, one from structural and one from functional brain features.

The BAG framework^50^ illustrated in Fig. 3a provides a compact subject-specific marker able to quantify individual deviations from expected brain values. The chronological age is predicted from either structural or functional brain features for each participant. For each participant, the BAG was defined as the difference between predicted brain age and chronological age after age-bias correction. Positive BAG indicates accelerated (higher-than-expected) brain aging, while negative BAG indicates delayed (lower-than-expected) brain aging^40,60–63^. Age prediction was computed by Ridge regression with 5-folds cross-validation. See Methods for details.

**Fig. 3:**
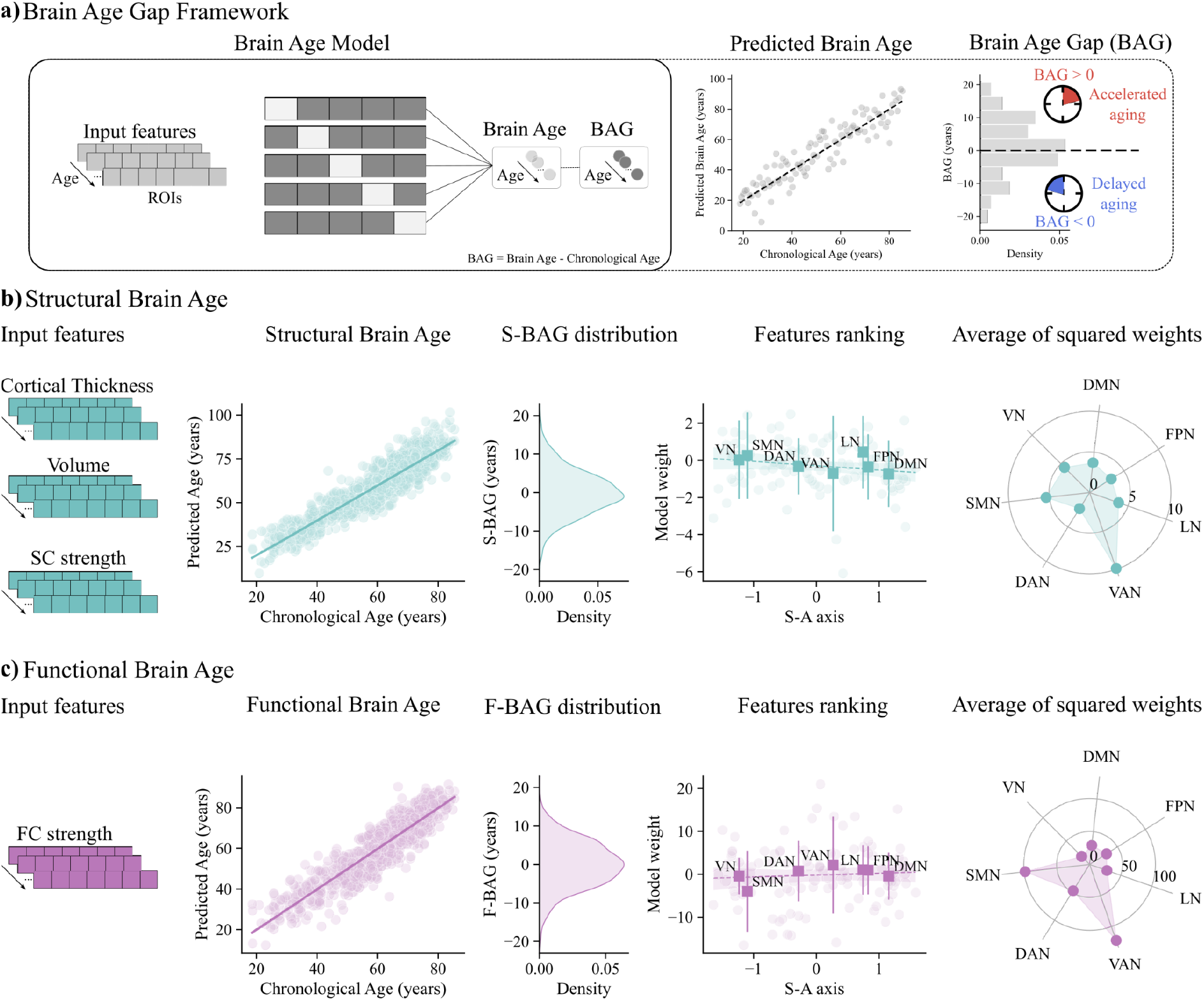
Structural and Functional Brain Age Models reveal distinct network signatures. **a)** Brain Age Gap (BAG) Framework. Participants’ ages are predicted from brain features through a 5-folds cross-validated approach. The regression residuals BAG = Predicted Age – Chronological Age can be positive or negative, thereby indicating higher or lower brain age compared to chronological age. **b)** Structural Brain Age. From left to right: input features (cortical thickness, volume, and SC strength combined), age-bias corrected structural brain age with respect to chronological age, structural BAG (S-BAG) distribution, model weights along the S-A axis (r = -0.17, p_spin_ = 0.06) with reported mean and standard deviation per RSN, average squared weights per RSN. **c)** Functional Brain Age. From left to right: input features (FC strength), age-bias corrected functional brain age with respect to chronological age, functional BAG (S-BAG) distribution, model weights along the S-A axis (r = 0.02, p_spin_ = 0.47) with reported mean and standard deviation per RSN, average squared weights per RSN.

#### Structural features predict age more accurately as compared to functional connectivity

We first built a Structural Brain Age Model, as shown in Fig. 3b, combining cortical thickness, volume and SC strength as regressors. This model reached performance corresponding to explained variance R^2^ = 0.71 (STD = 0.05 across folds) between predicted brain age and chronological age, similar to previously reported performances^64–66^. We then built a counterpart Functional Brain Age Model based on FC strength, as shown in Fig. 3c, which yielded an average R^2^ = 0.37 (STD = 0.05), similar to previous results based on functional connectivity^67,68^. Interestingly, S-BAG and F-BAG show similar distribution across subjects (Fig. 3b,c) with a standard deviation of 5.85 and 6.06 years respectively. For a fair comparison, note that cortical thickness, volume, and SC strength alone reached respectively R^2^ = 0.71, R^2^ = 0.71, and R^2^ = 0.53.

#### Structural and Functional Brain Age Models reveal distinct network signatures

We then analyzed the model weights associated with each ROI to interpret how the regression models combine the brain regions in order to predict the subjects’ age. Neither the structural nor the functional model weights were significantly associated with the S-A axis (Fig. 3b,c), suggesting that spatial organization of brain-age prediction did not follow the same cortical hierarchy previously observed for cognitive-profiles. Moreover, all RSNs contribute to the prediction both positively and negatively. To better characterize which RSNs were contributing the most to the prediction, we analysed the averaged squared weights per resting state network. This indeed revealed distinct network-level signatures across the two models, highlighting a not fully overlapping pattern of regional contribution (Fig. 3b,c). Both models showed the strongest contribution from VAN. For structural brain age prediction, VAN contribution is followed by SMN, VN, DAN and LN. Similarly, for functional brain age prediction, VAN is followed by SMN and DAN. Results suggest that structural and functional models differently capture whole-brain age-related changes, while attentional and unimodal regions tend to participate more in the prediction.

#### Structural and Functional BAG markers are positively associated

Although structural and functional brain age models differ in predictive accuracy and network-level signatures, we tested whether they were able to capture age-related deviations. To that end, we calculated the Spearman’s correlation across the population and found that S-BAG and F-BAG are positively associated (r = 0.21, p < 0.001; Fig. 4a). This indicates that individuals with an older-than-expected structural brain profile also tend to have an older-than-expected functional brain profile, and vice versa. To assess whether the association between the structural and functional BAG markers was influenced by age-range, we progressively higher the minimum age of the sample. When repeating the analysis across increasing age thresholds, the positive association slightly decreased, yet remaining significant (Fig. 4b). Thus, accelerated structural aging was partly reflected in accelerated functional aging, although the modest magnitude of the association suggests that the two markers could potentially capture distinct, possibly complementary, age-related information.

**Fig. 4:**
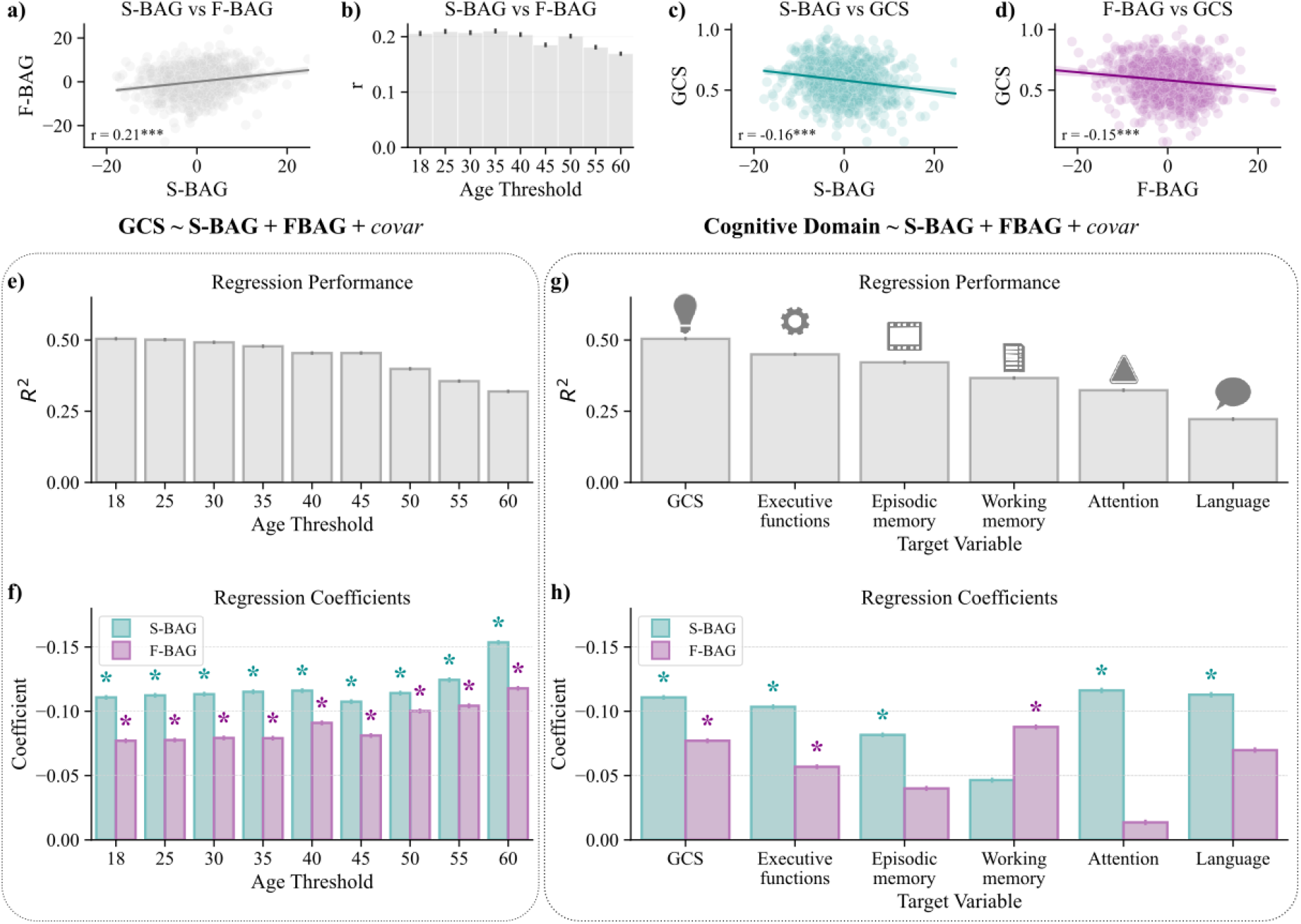
Structural and Functional brain age gaps as complementary and non-interchangeable markers of cognitive performance. **a)** Spearman’s correlation between S-BAG and F-BAG (r = 0.21, ***p < 0.001). **b)** Spearman’s correlation between S-BAG and F-BAG across age thresholds. **c)** Spearman’s correlation between S-BAG and GCS (r = -0.16, ***p < 0.001). **d)** Spearman’s correlation between F-BAG and GCS (r = -0.15, ***p < 0.001). **e)** GCS-targeted OLS regression model performance across age thresholds. **f)** S-BAG and F-BAG coefficients of GCS-targeted OLS regression across age thresholds. **g)** GCS- and domain-targeted OLS regression model performance on full sample. **h)** S-BAG and F-BAG coefficients for each OLS regression which target GCS and all cognitive domains on full sample. All error bars show the 95% CI across 1.000 subsamples (n = 400 subjects). * is present if the coefficient is considered to have an overall robust effect.

### Structural and Functional brain age gaps as complementary markers of cognitive performance

#### Global cognitive score is predicted by F-BAG as well as S-BAG

Given that S-BAG and F-BAG capture distinct aspects of age-related brain signatures, we next examined to which extent they inform about the subject’s cognition state, using GCS. As a first test reported in Fig. 4b,c,d S-BAG showed a slightly stronger statistical association with GCS (r = -0.16, p < 0.001) than F-BAG (r = -0.15, p < 0.001), but the similar magnitude for S-BAG and F-BAG suggests that both are associated with global cognition.

While previous studies investigated the association between cognition and brain age markers^35^, here we consider a single predictive framework to address the question whether the F-BAG provides additional information relevant to cognition when combined with S-BAG?

To that end, we predicted GCS as the target variable using S-BAG and F-BAG as regressor variables of interest, and age, age-squared, sex, and education as covariates. We employed ordinary least squares (OLS) regression with cross-validation (50% train, 50% test). See methods for details. Robustness was tested by repeating the GCS prediction with increasing minimum age (age threshold). For each age threshold, we subsampled 400 subjects 1.000 times (see Methods for details).

Regression performance decreases from an average R^2^ = 0.50 on the full age range, to R^2^ = 0.32 for minimum age of 60 years, shown in Fig. 4e. Decrease in accuracy may arise from higher inter-subject variability in late adulthood. To assess whether the low-dimensional BAG values were sufficient to capture cognitive variability, GCS was additionally predicted using the multivariate regression from cortical thickness, volume, SC strength and FC strengths using Ridge regression, as opposed to the univariate analysis presented in Fig. 2. Regression yielded a cross-validated R² = 0.47, similar to previous findings^20^, but slightly lower than the BAG-based model. Furthermore, when considering the BAG markers separately performance slightly decreases, suggesting that both markers are complementary contributing to the prediction (see Supplementary Section 5.1). This confirms that, despite the huge difference in dimensionality, the BAG markers are able to capture relevant information from neuroimaging measurements to robustly predict cognitive performance at the individual level.

As before, robustness check reported in Fig. 4f shows that S-BAG and F-BAG regression coefficients are stable and remain significant across all age thresholds (always while accounting for chronological age and covariates). Negative regression coefficients indicate older-than-expected structural (or functional) brain profiles tend to have poorer GCS, confirming expectations. Importantly, F-BAG contributes to a slightly lower but similar magnitude to the GCS prediction as compared with the S-BAG. Note that, among the covariates, education is the only covariate that is coherently significant across all age thresholds (see Supplementary Section 5.2).

#### Cognitive domains are heterogeneously predicted by brain age markers

Despite GCS being able to capture 40% of the cognitive tests variance, it is only a summary of the cognitive performance across all cognitive domains. To refine our investigation of how the BAG markers are related to cognition, we considered each cognitive domain separately. In this way, we ask whether S-BAG and F-BAG always jointly contribute to all domain prediction, or if they exhibit differences among the cognitive domains.

We thus repeated the OLS regression analysis for GCS and all cognitive domains: executive functions, episodic memory, working memory, attention and language. This results in six regression models to compare in terms of performances, but especially in terms of BAG markers contributions in order to predict the domain-specific performance.

In Fig. 4g the regression performances are shown for train and test sets across folds. Executive functions and episodic memory regressions show the highest prediction performances, followed by working memory, attention, and language. In Table 1 is reported the average performance across subsamples.

**Table 1:**
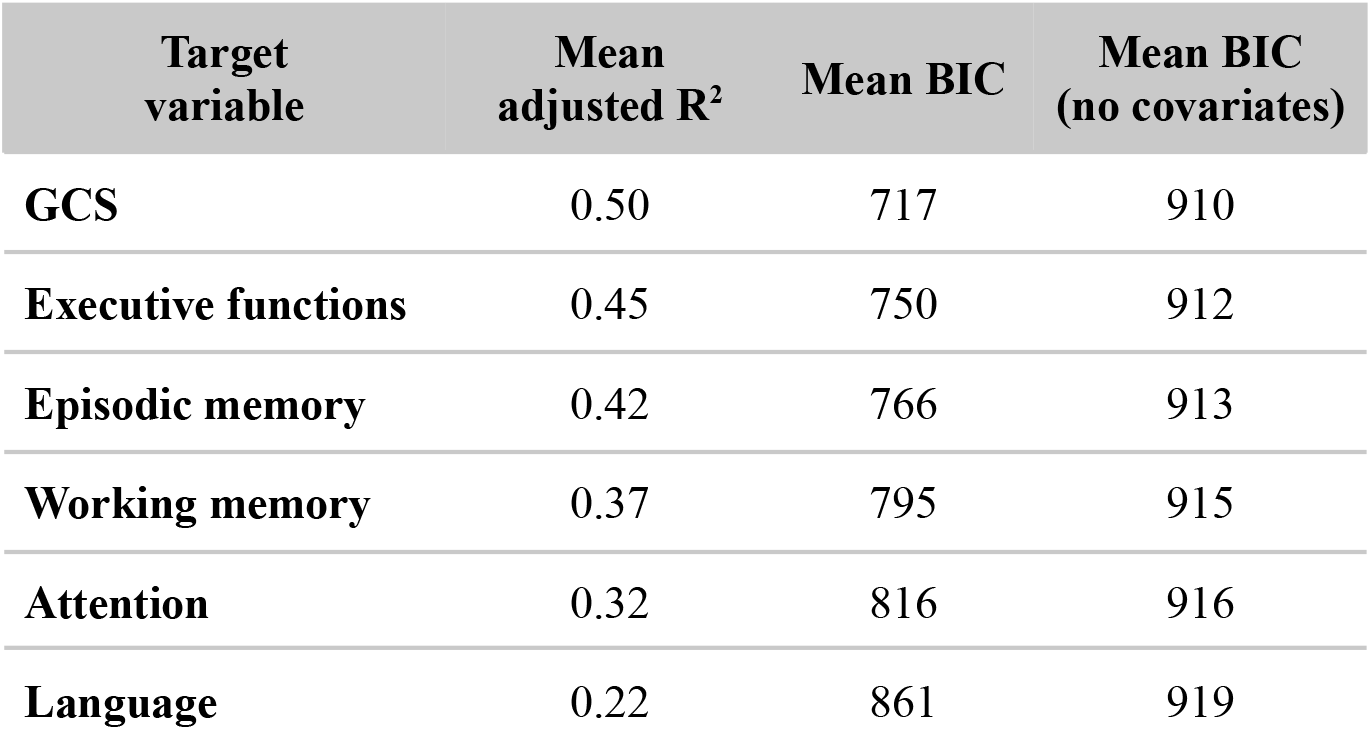
Average OLS regression performances across 1.000 subsamples.

Across all domains, both S-BAG and F-BAG model coefficients show negative contribution to the prediction, as shown in Fig. 4h and reported in Table 2. S-BAG shows robust contribution in predicting all domains, with exception for working memory. This is particularly interesting considering that working memory shows the highest contribution from the F-BAG marker. On the other hand, attention, episodic memory and language show greater contribution from S-BAG. Finally, it is especially striking that episodic and working memory are differentially associated with S-BAG and F-BAG. Overall, these findings suggest that structural and functional brain age deviations are not interchangeable, but specific and often complementary predictors of cognition.

**Table 2:** Average OLS regression coefficients across 1.000 subsamples. * is present if the coefficient is considered to have an overall robust effect.

| Target variable | S-BAG coefficient | F-BAG coefficient |
| --- | --- | --- |
| GCS | -0.11* | -0.08* |
| Executive functions | -0.10* | -0.06* |
| Episodic memory | -0.08* | -0.04 |
| Working memory | -0.05 | -0.09* |
| Attention | -0.12* | -0.01 |
| Language | -0.11* | -0.07 |

#### Parcellation robustness

To better assess robustness of results, we reproduced the BAG analysis on the 400 ROIs Schaefer parcellation. Robust accuracy in all cognitive domains was found, with slightly higher contribution of S-BAG and moderately lower contribution of F-BAG. The only discrepancies have been found in the domain-specific regression analysis. Particularly, the contribution of S-BAG for working memory, and the contribution of F-BAG for executive functions (see Supplementary Section 6). Nonetheless, while the two parcellations lead to similar age prediction accuracy, overfitting was greater using the 400 ROIs, coherently with previous benchmarks^69^.

## Discussion

How do brain structure and function respectively contribute to cognition? Here, we investigated how age-related patterns provide information about cognitive performance. By implementing a brain age gap (BAG) framework on the population-based 1000BRAINS cohort^55^, we extracted structural (S-BAG) and functional (F-BAG) markers, considering the deviation of the predicted brain age from the chronological age. We then demonstrated the added value of combining their contribution to explain global and domain-specific cognitive performance, providing a subject-level measure that can be compared across modalities.

### BAG markers as an efficient summary of cognitive-related brain information

We demonstrated that S-BAG and F-BAG provide a meaningful subject-level summary of relevant brain information for cognitive performance, with the two markers contributing to a comparable extent. This highlights that the accuracy in predicting chronological age from brain features does not guarantee a good performance for the corresponding BAG marker in predicting cognition and the corresponding inter-individual variability. Note that, in the extreme case of a perfect age prediction, the BAG as a regression residual would be zero for all subjects and thus uninformative. In our results, the structural brain age model predicted chronological age more accurately than the functional one, in line with previous evidence suggesting that structural measures more closely track biological aging^70^. However, both S-BAG and F-BAG have comparable contributions in predicting the cognitive domains, sometimes higher for F-BAG. Moreover, the regression to build the S-BAG relies on a distinct brain signature as compared to the direct prediction of cognition from structural phenotypes. This also holds for the F-BAG as compared to direct prediction of cognition from functional phenotypes. Neither BAG exhibits a statistical relation with the sensorimotor-association axis. Still, the contribution in age prediction of sensory and attentional regions to the functional model suggests that age-atypical functional deviations may involve lower-order systems, possibly reflecting changes in sensory processing, inhibitory control, or broader functional reconfiguration during aging.

### Structural and functional brain age gaps are complementary, yet not interchangeable

Structural and functional BAGs were only modestly correlated, indicating that structural and functional brain aging are partly coupled but appear to capture complementary dimensions of brain health. In fact, F-BAG contributed to the prediction of global cognition alongside the S-BAG, even after accounting for chronological age, age-squared, sex, and education. Thus, F-BAG still captured cognition-relevant inter-subject variability that was not fully explained by structural brain aging, meaning that structural integrity alone does not fully explain cognitive differences in normal aging.

### Structural and functional brain age gaps interplay in cognitive domains

Executive functions and episodic memory were the domains best predicted from the BAG markers, followed by working memory, attention, and language. This ordering is informative in itself. In fact, the BAG quantifies the deviation from the age-expected configuration, and therefore it is less informative where performance is less dependent on age. This pattern supports the validity of our framework.

#### Executive Functions

The executive functions domain was robustly predicted by both the S-BAG and F-BAG. This is in line with previous findings suggesting that decline is linked to poorer performance in domains requiring distributed coordination^71,72^, and that preserved structure–function integration of the salience/ventral attention system specifically tracks slower decline in executive performance in longitudinal cohorts^73^.

#### Episodic memory

The episodic memory domain includes tasks such as verbal list learning with delayed recall and figural retention. These rely on encoding and retrieving information, which depend on medial temporal and temporo-parietal integrity. Structural change in the hippocampus, amygdala and parahippocampal gyrus was the dominant correlate of episodic memory decline^74^, consistent with the brain maintenance account of successful memory aging^75^ and might explain why it was significantly predicted by the S-BAG only.

#### Working Memory

Working memory was the only domain for which F-BAG showed its strongest contribution. The domain consisted mainly of visuospatial tasks and verbal digit span. These tasks require maintaining and manipulating information encoded in sensory and attentional systems and their execution depends on a temporary reconfiguration of functional networks into a more integrated state, which correlates with individual performance^76–78^. Consistently, the capacity of a region to communicate broadly with the rest of the cortex is what predicts cognitive control performance^79^, rather than its local anatomy, in alignment with the F-BAG contribution.

#### Attention

The attention domain was assessed using the Trail Making Test A, which primarily measures processing speed, a key mediator of age-related differences across cognitive domains^80^ and closely linked to white matter integrity^81,82^. Since our structural model combined regional morphometry and structural connectivity, S-BAG likely reflects this substrate-related variability. Thus, aligning with the significant contribution of S-BAG.

#### Language

Similarly to episodic memory and attention, language was significantly predicted by S-BAG only, slightly in contrast with previous findings showing greater contribution from functional connectivity alone^83^. This could be due to different assessment tests, but also to sample biases such as mother tongue, and number of known languages, yet remaining the domain with the lowest prediction accuracy. This suggests that other hidden factors might be involved, especially considering the functional language network preservation in older adults^84^.

### Implications for Cognitive Aging Theories

A persistent challenge in cognitive aging research is how to characterize individuals who deviate from age-normative trajectories. Recent literature tried to overcome this issue by defining possible cognitive mechanisms, such as cognitive reserve, resilience, maintenance, and compensatory scaffolding^26,85,86^. Nonetheless, different cognitive outcomes despite similar age-related changes have been tested with heterogeneous operational definitions and indirect measures^87,88^. Superagers, for instance, have been identified using different thresholds, percentiles, or comparisons with younger reference groups^89^. These approaches are useful for isolating extreme groups, but they impose discrete boundaries, and classification depends on the chosen reference groups and cut-offs.

Our findings suggest that BAG markers may offer a complementary way to capture this variability, by providing individual-specific estimates of structural and functional deviation from age-expected brain patterns informative for cognitive performance. S-BAG and F-BAG might help characterize relative cognitive vulnerability and preservation without assigning individuals to predefined categories, in line with the idea behind normative modelling approach^90^. Lastly, the partly independent associations of S-BAG and F-BAG with cognition further suggest that structural, functional, and cognitive aging do not necessarily progress in parallel.

### Limitations and perspectives

Aging was inferred from differences between individuals rather than from within-subject trajectories. Longitudinal data will be required to determine whether the coupling between structural and functional BAG markers reflects progressive biological aging and compensatory functional reorganization^91^. Furthermore, recent work discussed that the brain age approach might index variations present since childhood^70,92,93^, nonetheless it may still serve to identify potential high-risk individuals. Besides, the functional brain age model provided relevant information for cognition, but functional connectivity remains sensitive to non-age-related sources of variability, including physiological noise, arousal, and state-of-mind during scanning. A further limitation concerns the features sets: the S-BAG model combined regional morphometry and connectivity, whereas the F-BAG model used a single measure per region, the nodal strength of the functional connectome. Further investigations could include other measures of intrinsic regional activity such as regional homogeneity or the fractional amplitude of low-frequency fluctuations. However this imbalance is unlikely to explain the difference in age-prediction accuracy, since cortical thickness and volume respectively reached R² = 0.71. Finally, future work is needed to investigate the identification of individuals at high risk for cognitive decline, by looking at subjects whose brain age significantly exceeds their chronological age across both modalities. Potentially, tracking the *biological* brain age longitudinally might be relevant for personalized medicine.

### Conclusions

In conclusion, the multimodal framework proposed here has demonstrated its potential in capturing structural and functional deviations with respect to the cognitive decline, shifting the paradigm from an age-prediction accuracy view to relevant subject-specific markers.

## Methods

### 1000BRAINScohort^17,18,21,47,55,94–97^

The population-based study is a subsample of the 10-year follow-up cohort of the Heinz Nixdorf Recall Study^98^. Non-eligibility for MRI measurements (due to e.g. coronary artery stents, surgical implants or prostheses in head or trunk, tattoos or permanent make-up on the head) was the only exclusion criterion. Each participant gave written informed consent prior to the admission in the cohort. The study protocol of 1000BRAINS was approved by the local Ethics Committee of the university Duisburg-Essen. Cohort comprises 1314 participants (44% females, 18-87 years, mean = 60.7, STD = 13.4), of which 901 subjects (43% females, 18-85 years, mean = 60.8 years, STD = 13.2 years) had all modalities and demographic information (see Supplementary Section 1 for age and education level distribution).

### Imaging data acquisition and preprocessing

Imaging data were acquired on a 3T Siemens Tim-Trio MR scanner (32-channel head coil; Siemens, Erlangen, Germany) at the Forschungszentrum Jülich, Germany. High-resolution T1-weighted anatomical images were obtained using a 3D MPRAGE sequence (TR = 2250 ms, TE = 3.03 ms, flip angle = 9°, voxel size = 1 mm³). Resting-state functional images were acquired with a T2*-weighted gradient-echo EPI sequence (TR = 2200 ms, TE = 30 ms, voxel size = 3.1 mm³, 300 volumes, scan duration ≈ 11 min). Participants were instructed to keep their eyes closed, remain awake, and let their minds wander freely without engaging in any structured cognitive task. Two diffusion datasets with two different b-values were acquired, for subsequent multi-shell fiber tract reconstruction, using the following sequence parameters: b=1000 s/mm² (EPI, TR=6.3 s, TE=81 ms, 7 b0-images (interleaved) and 60 diffusion-weighted volumes) as well as b=2700 s/mm² (EPI, TR=8 s, TE=112 ms, 13 b0-images (interleaved) and 120 diffusion-weighted volumes), both at 2.4 mm isotropic resolution.

Structural T1-weighted images were processed with the automated surface-based pipeline implemented in FreeSurfer^99^ to derive regional cortical thickness and volume estimates. High-angular resolution diffusion imaging (HARDI) was acquired to enable tractography-based reconstruction of white matter fiber pathways, from which structural connectivity matrices were derived. Main pre-processing steps included the removal of Gibbs, eddy currents and motion artifacts from the data-sets, correction of susceptibility-induced distortions, normalization of signal intensities as well as spatial correction of distortions between the images acquired at different b-values. Fiber tract reconstructions were based on the local constrained multi-shell, multi-tissue spherical deconvolution modelling^100^ of the diffusion signal as implemented in MRtrix3^101^ on the integrated and masked diffusion data. 10 million streamlines were generated seeding from the grey-white matter interface using the iFOD2 algorithm in MRtrix. The connectivity for each region*region combination was estimated as streamline counts for each participant.

Resting-state BOLD time series were preprocessed using the standard fMRIPrep^102^ pipeline, applying a low-pass filter at 0.1 Hz. Global signal regression was not applied. All modalities were parcellated according to the 100 and 400 regions of interest (ROIs) Schaefer atlas^57^, with regions further grouped into the seven canonical resting-state networks defined by Yeo^58^.: visual (VN), somatomotor (SMN), dorsal attention (DAN), ventral attention (VAN), limbic (LN), fronto-parietal (FPN), and default mode (DMN). The Schaefer parcellation with 400 regions was used for replication purposes.

### Functional connectivity and nodal strength

Functional connectivity was computed from the BOLD time series as the Fisher’s z-transformation of the Pearson’s correlation between the activation of pairs of regions, resulting in a symmetric connectivity matrix NxN ROIs. Additionally, for the connectivity-based measures (SC and FC), we computed the nodal strength for each region of interest, by summing all the incoming connections from all other regions, resulting in a nodal vector of N ROIs.

### Cognitive assessment

Together with the imaging recordings, a battery of neuro-psychological assessment was conducted. A total of 13 cognitive tests were considered, spanning different cognitive domains such as executive functions, attention, language, episodic and working memory (for a detailed description^17,94,103^). In order to keep as many participants as possible, NaNs values were replaced with the average score for the matched age range (5 years above and below). We excluded all subjects with more than 20% missing values. In order to investigate the population-based cognitive decline and link it to the brain aging process, we performed a Principal Component Analysis (PCA) on the 13-dimensional cognitive space. The first principal component, here called Global Cognitive Score (GCS) was found to explain 40% of the total variance. To characterize the global tendency across the lifespan, we then fitted a polynomial regression. The quadratic regression was found to best explain the trajectory of the GCS. Based on the subject-specific residual from the predicted mean, we artificially stratified the subjects using the 20th and 80th percentiles, in order to compare discrete cognitive profiles of “best performers” versus “poorest performers” in Fig. 1.

#### Brain Age Framework

Chronological age was predicted from the feature set of brain phenotypes using a supervised regression framework^38^, as follows. Structural brain age was predicted from the combination of cortical thickness, volume and SC strength, for a total of 300 features on a sample of 1212 participants (43% females, 18-85 years, mean = 60.8 years, STD = 13.3 years). Functional brain age was predicted from FC strength only, for a total of 100 features on a sample of 925 participants (43% females, 18-85 years, mean = 60.8 years, STD = 13.1 years).

##### Brain Age Models

The pipeline consists of a nested-cross validation scheme, implemented in Python (v3.13) using the scikit-learn^104^ toolbox. The outer cross-validation loop consisted of stratified k-fold splitting (k = 5), where stratification was performed using quantile-based bins of the target variable to preserve the age distribution across folds. For each outer fold, the training set was used for model selection, while the test set served exclusively for performance evaluation. Within each outer training set, hyper-parameters were optimized using a randomized search procedure sampling from predefined distributions. Hyper-parameter search was performed using a stratified 5-fold cross-validation scheme. The model achieving the best cross-validated score (R² unless otherwise specified) in the inner loop was selected and refitted on the full outer training set. The optimized model was then used to aggregate predictions for each test set fold to obtain out-of-sample predictions for all participants.

##### Model selection

Model performance was quantified using multiple metrics, including coefficient of determination (R²), mean absolute error (MAE), root mean squared error (RMSE), and Pearson’s correlation between predicted and true chronological age. Metrics were computed separately across folds for training and test sets. Training targets were randomly shuffled and refitted to obtain a chance-level performance within each outer fold (1.000 permutations).

We benchmarked our BAG pipeline against the AgeML^38^ toolbox across different regressors, scaling methods and scoring types. In particular, we considered the Ridge, Lasso and Support Vector (SVR) regressions, with scoring as MAE, RMSE and R^2^. As scalers we tested the Standard, MinMax and Robust. As shown in Fig. 5 our BAG pipeline exceeded the AgeML toolbox in terms of accuracy and overfitting, for both structural and functional models. The selected model, achieving best R^2^ on the test set was a MinMax scaler, Ridge regression with MAE scoring strategy (see Supplementary Section 4.2-4.4 for all performance metrics and sensitivity analysis).

**Fig. 5:**
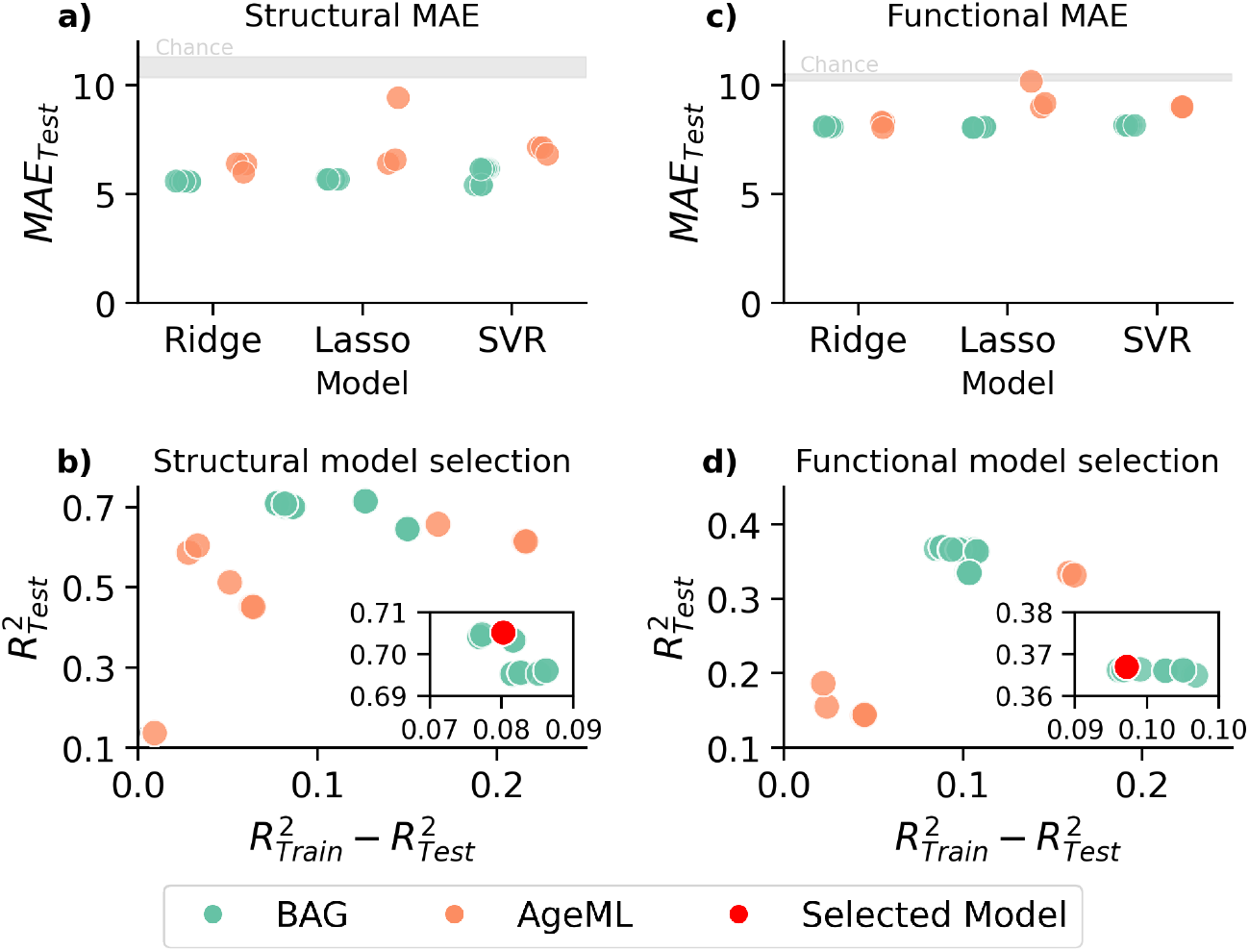
Brain age model selection. **a)** MAE of test set for the different regression models. Our BAG pipeline (green)is benchmarked against the AgeML (orange) for age prediction from cortical thickness, volume, and SC strength combined. **b)** Structural brain age model selection. R^2^ of the test set is shown again the overfitting (train - test). Zoom shows the models with highest test-set accuracy and the selected model (red). **c)** MAE of test set for the different regression models. Our BAG pipeline (green) is benchmarked against AgeML (orange) for age prediction from FC strength. **d)** Functional brain age model selection. R^2^ of the test set is shown again the overfitting (train - test). Zoom shows the models with highest test-set accuracy and the selected model (red).

##### Age-bias correction

To overcome the regression to the mean problem^105^, we then applied the age-bias correction on the model predictions. Let *Y* be the chronological age and *Y*’ the predicted age. We define the brain age gap as the difference between the predicted and real age, meaning *BAG* = *Y*’ − *Y*. We initially fitted the linear model *BAG* = α * *Y* + β, to then compute the residual brain age gap, i.e. *correctedBAG* = *BAG* − (α * *Y* + β). We derived the age-bias corrected prediction as *corrected Y*’ = *correctedBAG* + *Y* (see Supplementary Section 4.5 for age-bias correction influence on predictions).

#### Statistical Analyses

##### Group-level analysis

All statistical analyses were performed with a permutation-based statistics, using the FDR correction^106^ for adjusting for multiple comparisons (α = 0. 05 for significant effects). All reported p-values are corrected. Effect sizes are obtained using the rank-biserial index (*rb*)^107^, which is considered to express in absolute value a small effect if *rb* ≈ 0. 10, a medium effect if *rb* ≈ 0. 30, and a large effect if *rb* ≥ 0. 50^106^.

##### Brain maps correlation analysis

To assess whether two cortical maps were associated, we used a spatial permutation test, also known as spin test, to account for the spatial autocorrelation, which can inflate false-positive rates under standard parametric testing. In particular, we considered the Spearman’s correlation between two cortical maps of the same length, i.e. both maps have the same number of regions. Null distributions were generated by applying random rotations to the 3D parcel centroid coordinates, independently mirrored across hemispheres to preserve anatomical symmetry, and reassigning each parcel the value of its nearest rotated neighbor^108–110^. This procedure was repeated 1,000 times to build the null distribution of correlations for each comparison. p_spin_ were calculated as the proportion of null correlations at least as extreme as the observed value, using the ENIGMA^109^ toolbox.

##### Multivariate regression analysis

Independent contributions of structural and functional brain ages to cognitive variability were assessed using multivariate regression models with the global cognitive score as target variable. The main OLS regression included both the S-BAG, and F-BAG as predictors, adding as covariate the chronological age and its quadratic term, together with sex and education, thereby allowing the unique contribution of each BAG measure to be estimated while accounting for non-linear age effects. Note that here age-bias correction does not affect the regression accuracy, since it is based on a linear model.

To test whether these subject-level BAG indices captured information comparable to that contained in the original imaging feature space, a second model predicted the same cognitive outcome from age, age squared, and the full set of regional structural as well as structural and functional connectivity strength measures. Because this second model involved a high-dimensional predictor space, it was implemented using Ridge regression (see Supplementary Section 5.3 for details).

Analyses were repeated using an increasing minimum age threshold to evaluate whether the relative contribution of summary BAG measures and full regional features remained stable across adulthood. Since sample size decreases across age thresholds, we drew 1.000 subsamples of 400 subjects (without replacement), stratified jointly by age quintile and sex to preserve the original age-sex distribution. Within each subsample, regression model performance was quantified using stratified 5-fold cross-validation with age-binned splits, and out-of-sample R² was used for model comparison. Each independent variable’s coefficient was averaged across folds, yielding one coefficient per subsample. A variable contribution at a given age threshold was considered robust if the 95% percentile interval of its coefficient distribution excluded zero. All Multivariate regression analysis were implemented using statsmodels^111^ and scikit-learn^104^.

## Data Availability

Access to neuroimaging and phenotypic data is restricted due to ethical and data protection regulations. Data is available to qualified researchers upon reasonable request to the 1000BRAINS study organizers^55^.

## Code Availability

Code for extracting the brain age gaps is available at https://github.com/brainets/bag_multimodal. All analyses were performed in Python (v3.13) using numpy^112^, pandas^113^, statsmodels^111^, scikit-learn^104^ and ENIGMA^109^.

## Acknowledgements

Authors acknowledge the PEPR Santé Numerique for funding the research. This project was partially funded by the 1000BRAINS study of the Institute of Neuroscience and Medicine, Research Center Juelich, Germany. We thank the scientists and the study staff of the Heinz Nixdorf Recall Study and 1000BRAINS. The authors are supported by the European Union’s Horizon Europe Program under the Specific Grant Agreement No. 101147319 (EBRAINS 2.0 Project; SC). We gratefully acknowledge the computing time granted through JARA-HPC on the supercomputer JURECA at Research Centre Jülich (Germany). MG received support from the French government under the France 2030 investment plan, under the agreement Chaire de Professeur Junior (ANR-22-CPJ2-0020-01) and as part of the Initiative d’Excellence d’Aix-Marseille Université -A*MIDEX (AMX-22-CPJ-01).

## Ethics declarations

We declare no competing interests.

## Supplementary Information

SupplementaryInformation.pdf

