## Supplementary Information for "Functional and Structural Brain Age Gaps as Complementary Predictive Markers for Cognitive Performance"

#### 1. 1000BRAINS population and education level

The age distribution of the 1000BRAINS cohort is shown in Fig. S1, together with the education level, both stratified by sex. Here we considered the sample of participants with all brain modalities and demographic data, composed of 901 participants (43% females, 18-85 years, mean = 60.8 years, STD = 13.2 years). Education is referred to in the International Standard Classification of Education (ISCED 1997).

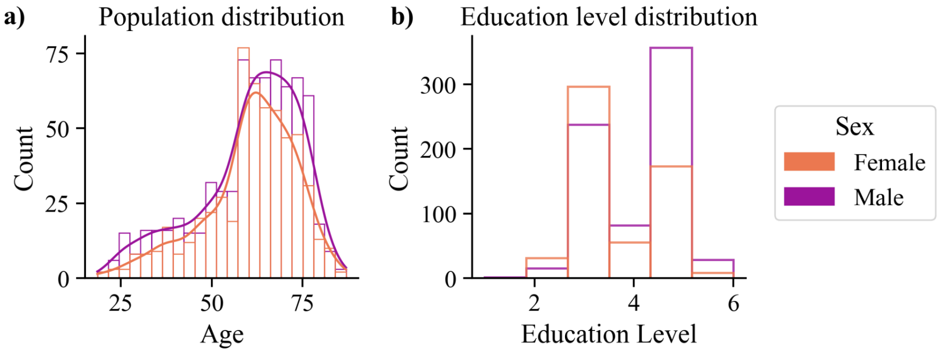

**Figure S1: 1000BRAINS population. a)** Age distribution stratified by sex. **b)** Education level distribution stratified by sex. Education is referred to in the International Standard Classification of Education (ISCED 1997). Levels are the following: lower secondary education (2), upper secondary education (3), post-secondary (non tertiary) education (4), first step of tertiary education (5), second step of tertiary education (6).

#### 2. PCA on cognitive scores

Here we report the specifics on the Principal Component Analysis (PCA) performed on the full set of cognitive scores, in order to extract a global cognitive score (GCS), and a domain-specific score for the following domains: attention, executive functions, language, working memory, and episodic memory.
In Fig. S2.1 the scree plot of the PCA analysis on all scores for the first 10 components with the respective explained variance ratio is shown. The first principal component (PC) explains 40% of the total variance. Fig. S2.2 shows all PCA loadings for the first 5 PCs. First PC shows similar loadings from all scores, with problem solving and figural memory being the slightly highest contributors.
Finally, in Fig. S2.3 the GCS and all the firsts PCs of the domain-specific PCA along the age axis are shown. All extracted components report significant negative correlation with age. All extracted components are normalized between 0 and 1 to facilitate interpretation. All values of explained variance for all domains and correlations with age are reported in Table S2.

| Domain | Explained Variance | Spearman's r with age | FDR corrected p value |
|---|---|---|---|
| **GCS** | 40.24% | -0.65 | ***p<0.001 |
| **Episodic memory** | 72.06% | -0.65 | ***p<0.001 |
| **Executive functions** | 58.90% | -0.62 | ***p<0.001 |
| **Working memory** | 49.96% | -0.53 | ***p<0.001 |
| **Attention** | 100% | -0.62 | ***p<0.001 |
| **Language** | 59.20% | -0.34 | ***p<0.001 |

**Table S2**: Cognitive principal components with their respective explained variance, Spearman's correlation with age, and corrected associated values.

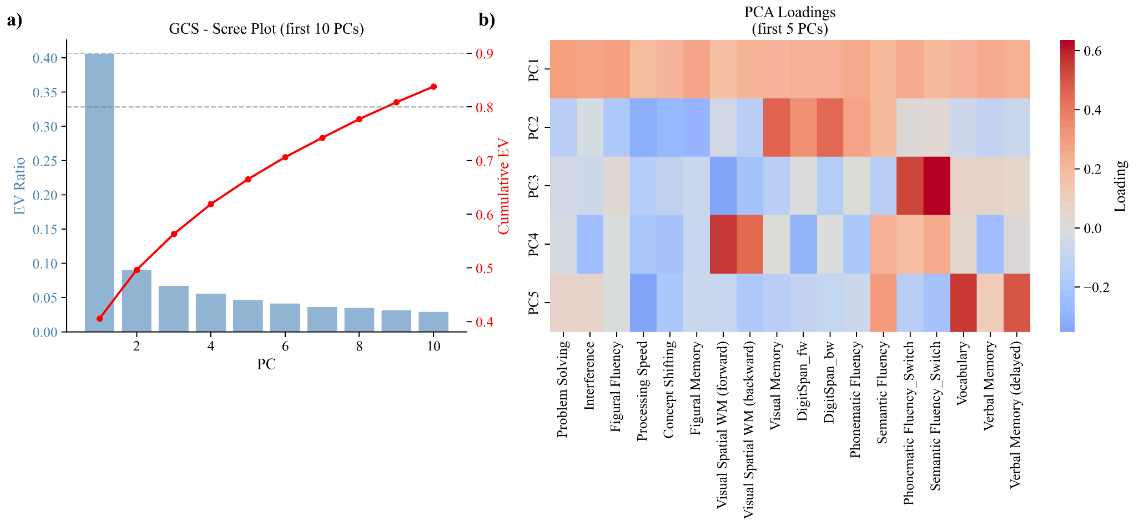

**Figure S2.1: PCA on the full set of cognitive scores. a)** Scree plot for the first 10 PCs. b) Heatmap of the PCA loadings for the first 5 PCs. PC1 is the one we considered as the global cognitive score (GCS) in the main text.

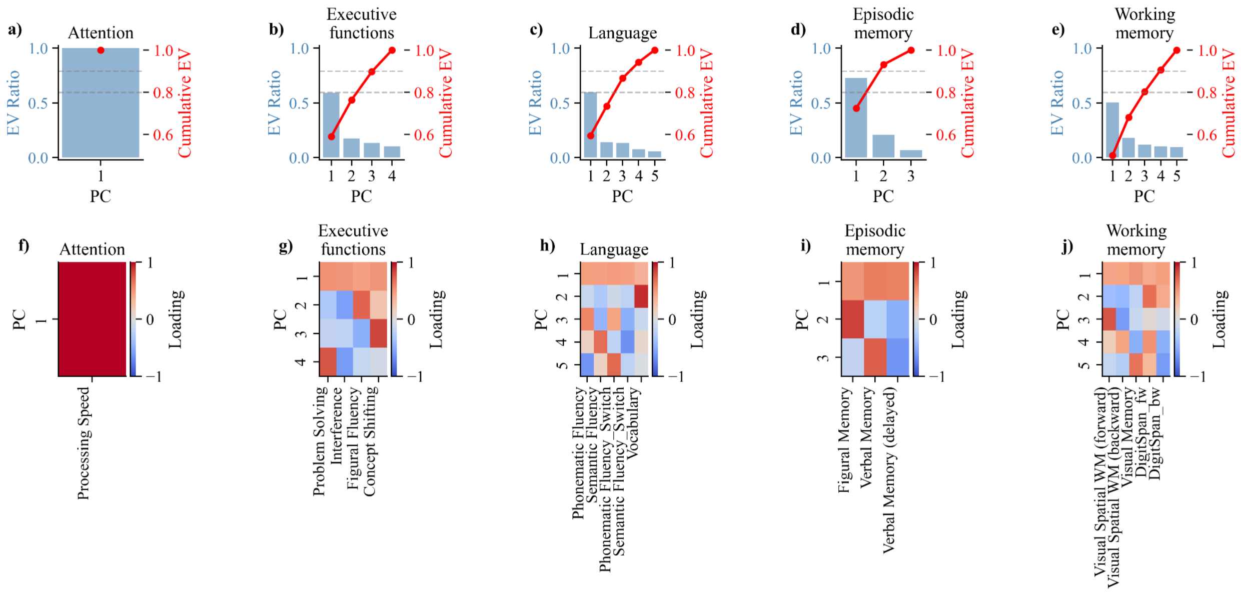

**Figure S2.2: Domain-specific PCA. a)-e)** Scree plot for all cognitive domains. **f)-j)** PCA loading for all scores in the cognitive domain.

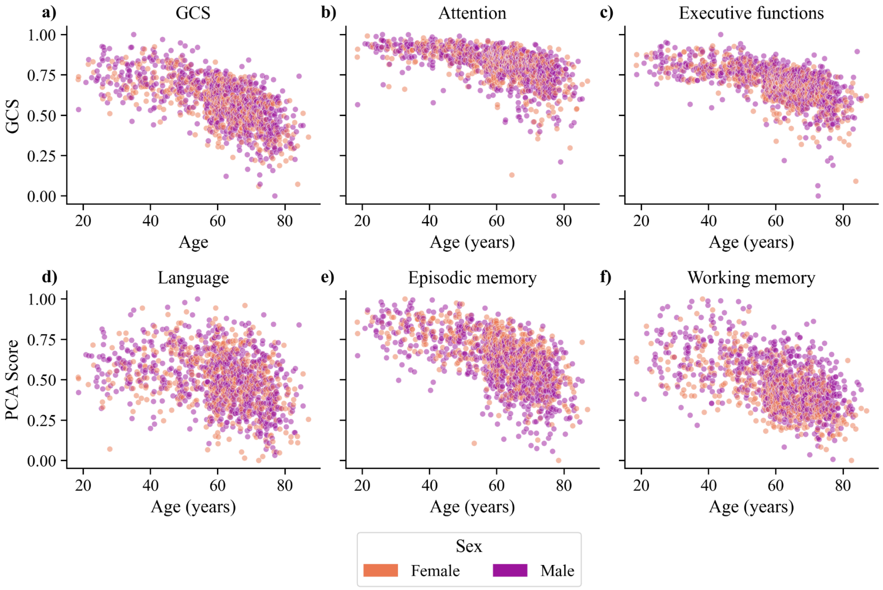

**Figure S2.3: GCS and domain-specific PC along the age axis, stratified by sex. a)** Global Cognitive Score (GCS). **b)** Attention. **c)** Executive functions. **d)** Language. **e)** Episodic memory. **f)** Working memory.

### 3. Cognitive profiles stratification analysis

#### 1. Age bins stratification

Since the cognitive profiles average analysis is compressing all the information related to age, we considered age-bins in order to see if age-related changes could indeed create a bias in the interpretation. As shown in Fig. S3.1, the young-adult bin (18-40 years) is showing a slightly different pattern from the adult and elder bins. In fact, in young adults the difference in FC strength seems to be stronger, mostly due to higher FC strength in the high cognitive group.

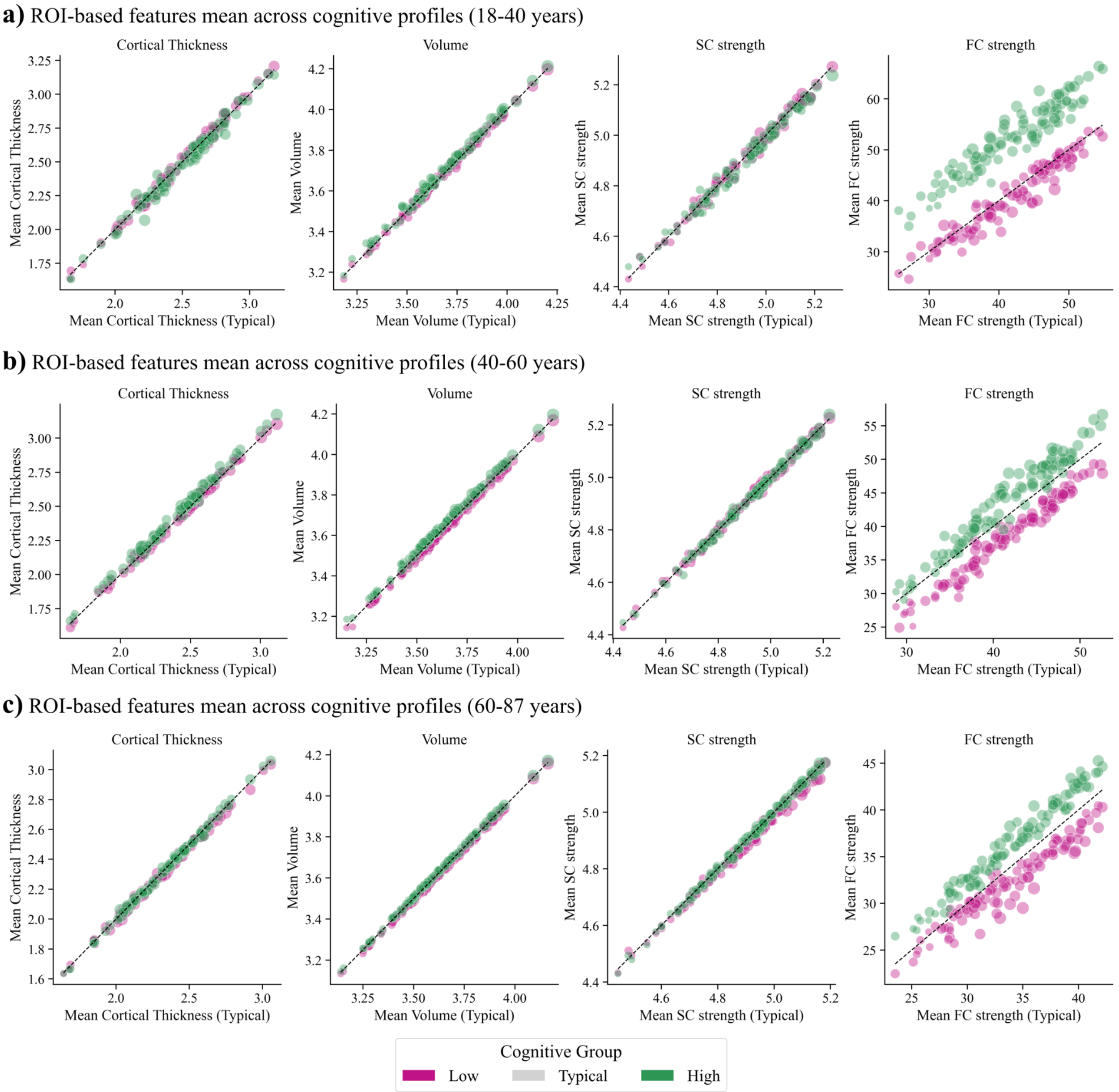

**Figure S3.1: ROI-based features across cognitive groups and age bins.** Each dot represents one ROI. The values represent the mean values of the high (green) or low (pink) with respect to the typical cognitive group (dashed line). **a)** Young-adults (18-40 years). **b)** Adults (40-60 years). **c)** Elders (60-87 years).

#### 2. Sex stratification

In order to see whether in the brain features we were able to see differences related to sex, we also repeated the cognitive profiles analysis stratifying participants based on sex. As shown in Fig. S3.2, no sex differences are found in high and low cognitive profiles for FC strength. On the other hand, standard deviation values across brain features and ROIs reveals a sex difference, with more heterogeneous differences in standard deviation in females across the ROIs.

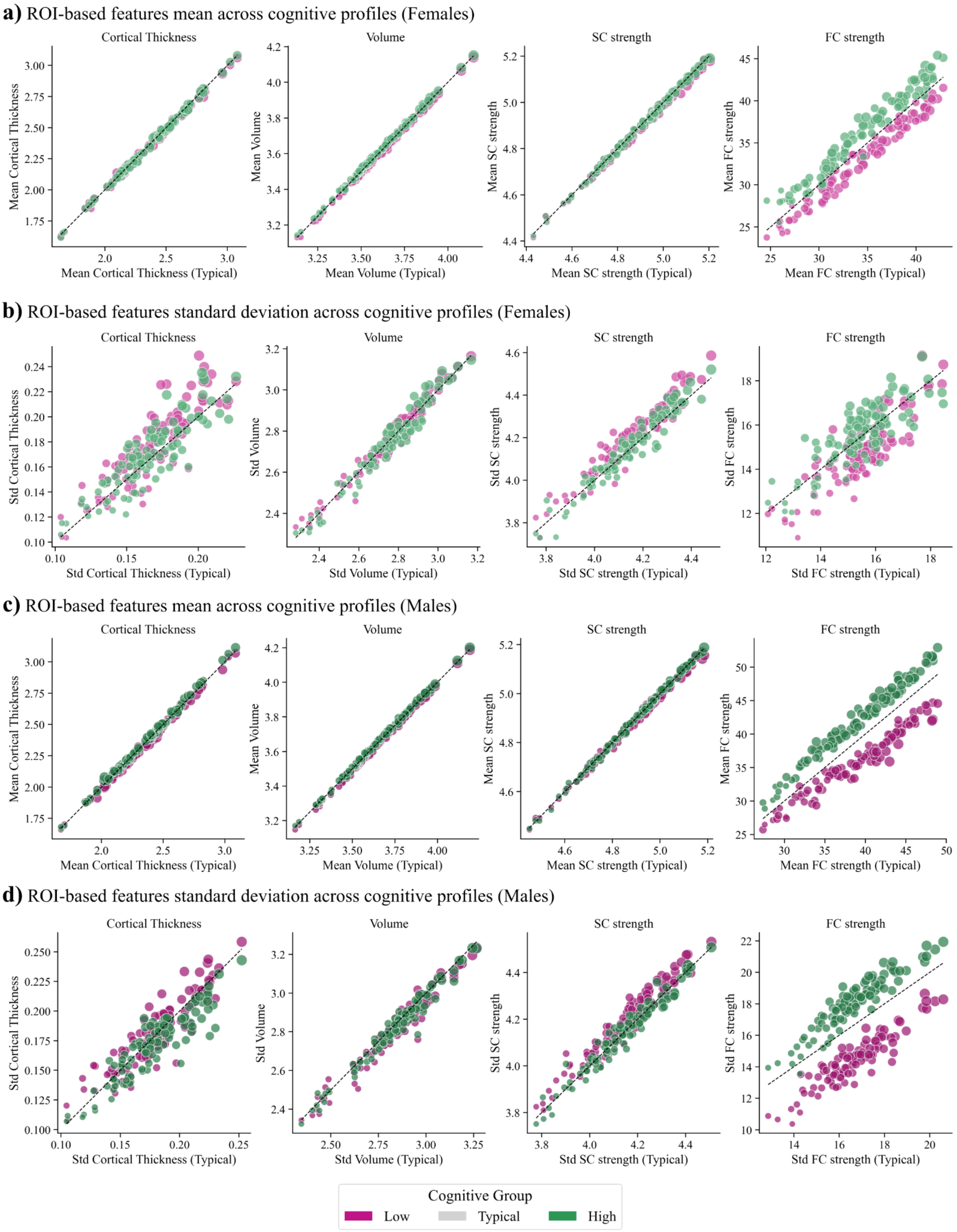

**Figure S3.2: ROI-based features across cognitive groups divided by sex. Each dot represents one ROI. a)** Mean values in females. **b)** Standard deviation values in females. **c)** Mean values in males. **d)** Standard deviation values in males.

### 4. Brain Age Models

To ensure that the brain age estimates were not driven by an arbitrary modelling choice, we performed a series of supplementary robustness analyses. First, we compared different parcellation resolutions and feature representations, including nodal-strength and full-matrix models. Second, we benchmarked our selected pipeline against the AgeML toolbox. Third, we assessed the alignment of BAG estimates across alternative models by quantifying BAG differences, variability across models, and sign agreement. These analyses supported the use of the Schaefer 100-ROI nodal-strength representation combined with the selected Ridge-based pipeline for the main analyses.

#### 1. Parcellation selection

To select the parcellation to use in brain age predictions we first tested different input features. Here we compare three different combinations.

For the structural model we compared the following input features:

- 300 features for parcellation N = 100 strength: cortical thickness, volume and SC strength in 100 ROIs parcellation,
- N = 100 full: cortical thickness, volume and SC upper triangular matrix (4950 features), for a total of 5150 features
- N = 400 strength: cortical thickness, volume and SC strength in 400 ROIs parcellation

For the functional model we compared the following input features:

- N = 100 strength: FC strength in 100 ROIs parcellation,
- N = 100 full: FC upper triangular matrix, corresponding to 4950 features (or pairwise FC edges)
- N = 400 strength: FC strength in 400 ROIs parcellation

In Fig. S4.1 are reported the performances across folds of each combination of input features for both structural and functional models.

We selected as features the 100 ROIs parcellation, using the nodal strengths instead of the matrices for two main reasons: computational cost is reduced when the number of input features is lower, and in both structural and functional models it was the combination with the lowest overfitting (difference between accuracy in train and test set).

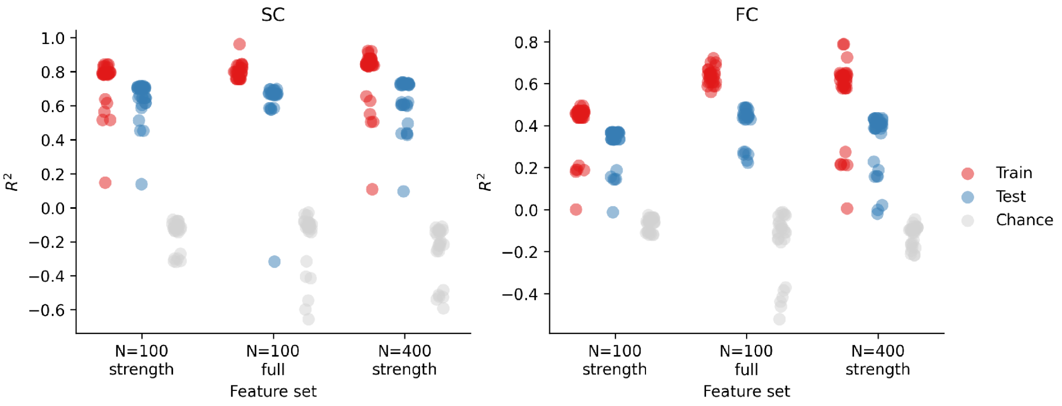

**Figure S4.1: Parcellation performance comparison. Left:** structural brain age model performances across different input features. Right: functional brain age model performances across different input features.

##### 2. Structural models performances

We tested our pipeline against the open source AgeML toolbox. All performance metrics are reported in Fig. S4.2 for structural features.

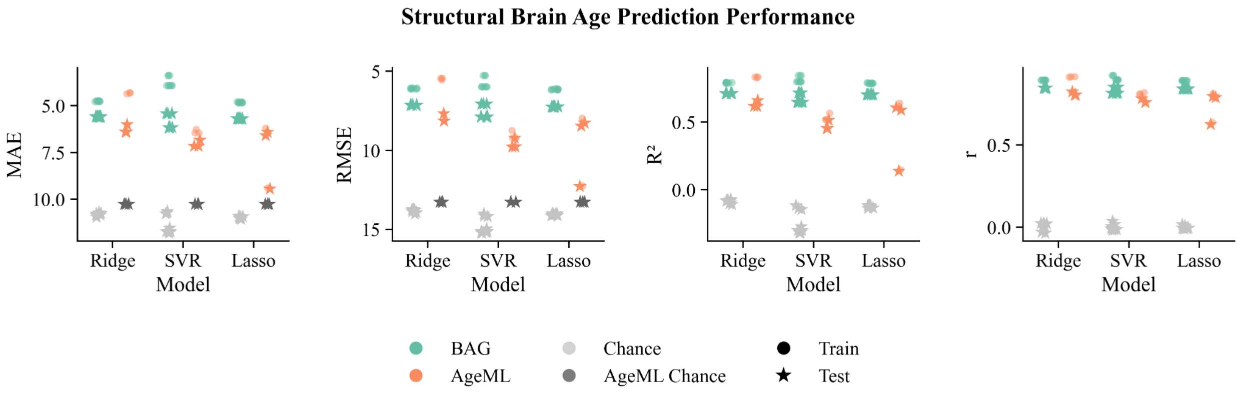

**Figure S4.2: Structural models performance.** Chronological age is predicted from cortical thickness, volume and SC strength combined.

##### 3. Functional models performances

We tested our pipeline against the open source AgeML toolbox. All performance metrics are reported in Fig. S4.3 for functional features.

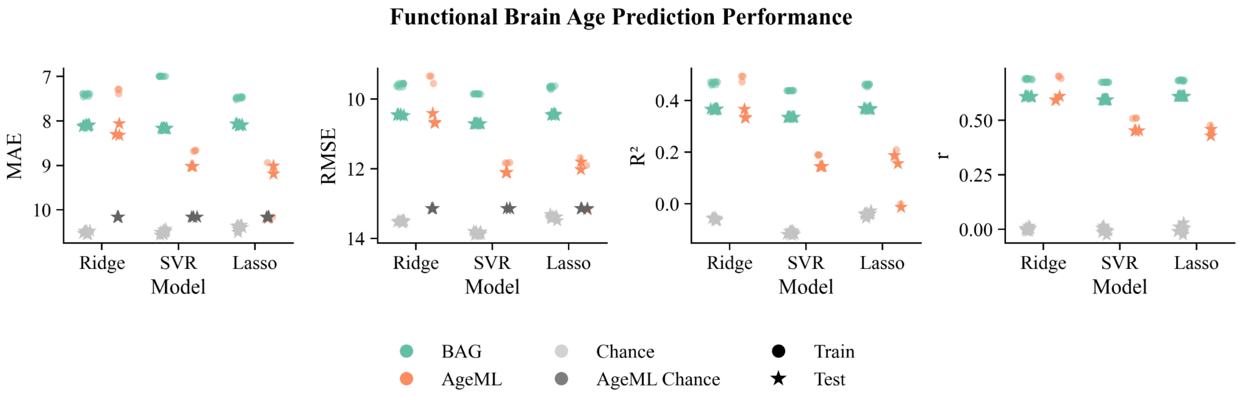

**Figure S4.3: Functional models performances.** Chronological age is predicted from FC strength. Left: $R^2$ between predicted age and chronological age in train and test sets.

##### 4. Sensitivity analysis across models

To assess whether age predictions were consistent across models, we considered the difference between the predicted BAG for two models across subjects. The average difference between BAG tests the amplitude of differences between predictions across subjects. The standard deviation of BAG differences tests the reliability across models. Furthermore, we are interested in understanding whether participants have coherent sign prediction across models, meaning whether participant's brain features predict higher-than-expected age in all models. To investigate this we use the percentage of sign agreement between two models.

Overall agreement between models is high, meaning that brain age prediction and the corresponding BAGs are replicable with different models. Interestingly, the functional brain age model shows higher sign agreement and lower BAG differences across models with respect to the structural brain age model (see Fig. S4.4), suggesting that functional models have more coherent predictions across different pipelines.

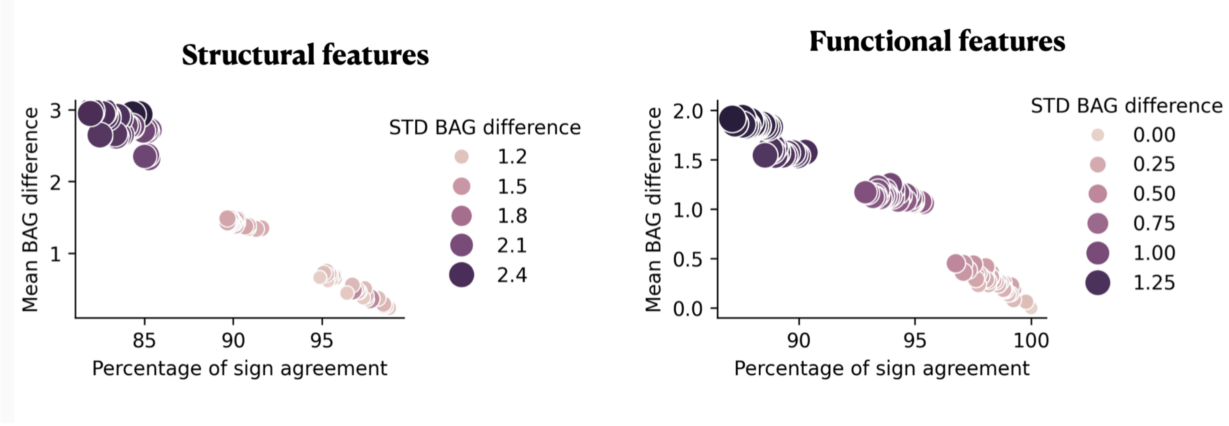

**Figure S4.4: Sensitivity analysis across models.** Each dot represents the comparison between a pair of models.

##### 5. Age-bias correction influence on predictions

When using the Brain Age Models approach to predict participants' age, a common problem is the so-called regression to the mean problem, for which ages at the extreme of the age range tend to be predicted closer to the average age. To overcome this issue, model predictions are then corrected. This phenomenon is clearly visible in Fig. S4.5, especially in the case of functional brain age predictions.

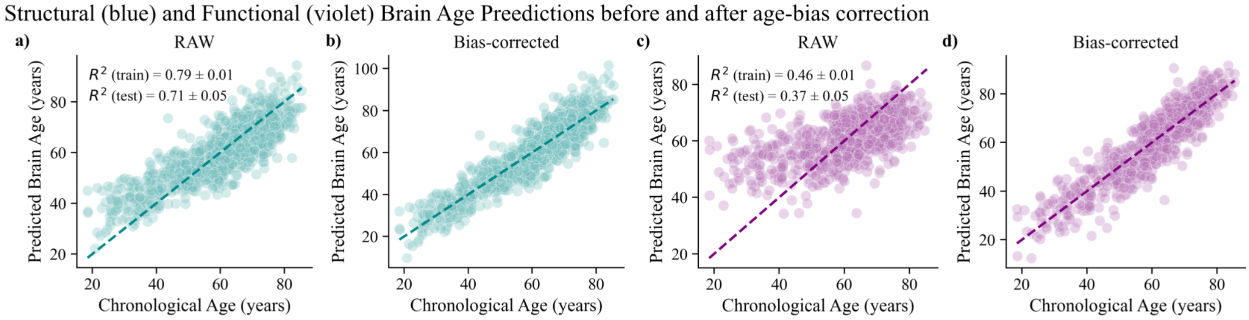

**Figure S4.5: Comparison of brain age predictions before and after age-bias correction. a)** Raw and **b)** age-bias corrected Structural Brain Age predictions. **c)** Raw and **d)** age-bias corrected Functional Brain Age predicitons.

##### 6. BAG markers correlation with GCS

Here we investigate further the correlation between the BAG markers and the GCS, by increasing the minimum age of the considered sample. As it can be seen in Fig. S4.6, both for S-BAG and F-BAG, correlation increases when increasing the minimum age. To better understand whether the changes across age thresholds were driven by age effects or sample size, we subsampled 400 subjects for each age threshold 1000 times. Notably, even after matching sample sizes, we could still see some age effect in the correlation between structural and functional BAGs, and between BAGs and GCS.

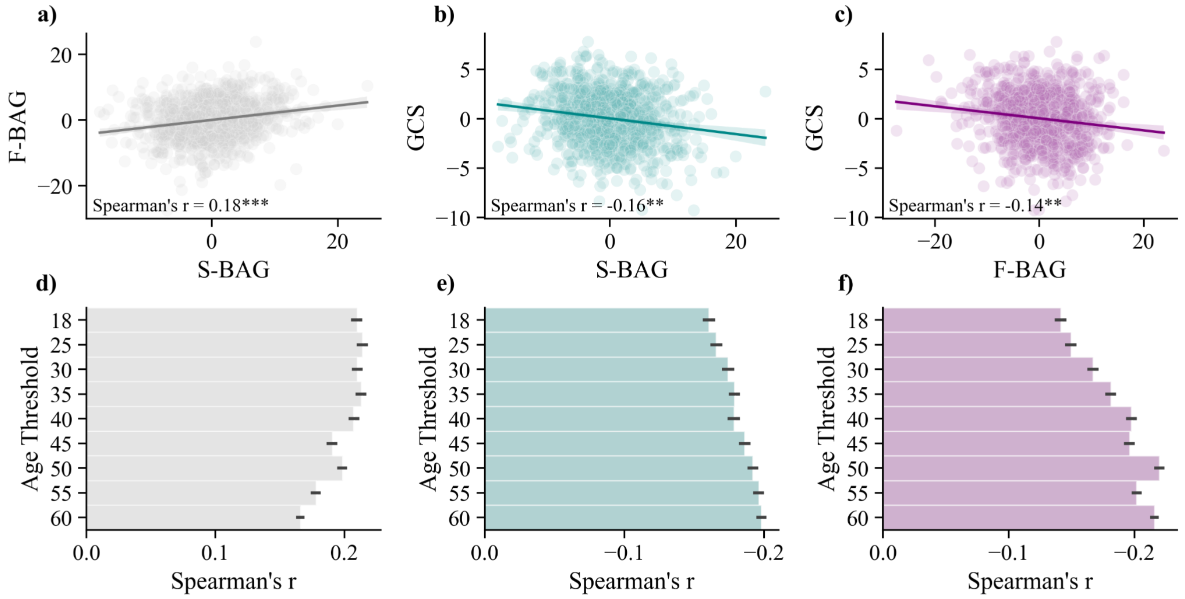

**Figure S4.6: BAG markers correlation analysis. a)** Correlation between S-BAG and F-BAG on full sample. **b)** Correlation between S-BAG and GCS on full sample. **c)** Correlation between F-BAG and GCS on full sample. **d)** Subsampling correlation analysis between S-BAG and F-BAG across age thresholds. **e)** Subsampling correlation analysis between S-BAG and GCS across age thresholds. **f)** Subsampling correlation analysis between F-BAG and GCS across age thresholds. Subsampling was performed randomly sampling without replacement 400 subjects for each age threshold (1000 times). Age distribution and sex ratio was maintained in the sampling.

#### 5. Multivariate regression models for global cognition

##### 1. S-BAG and F-BAG separate contribution to GCS

We tested whether S-BAG and F-BAG separately could inform GCS regression. Fig. S5.1 shows the adjusted $R^2$ across age thresholds and subsamples for the three proposed models: S-BAG + F-BAG, S-BAG, and F-BAG. For all age thresholds, the model with both structural and functional BAG markers shows higher $R^2$, suggesting that taken together, structure and function might give a more refined information about the subjects' cognition.

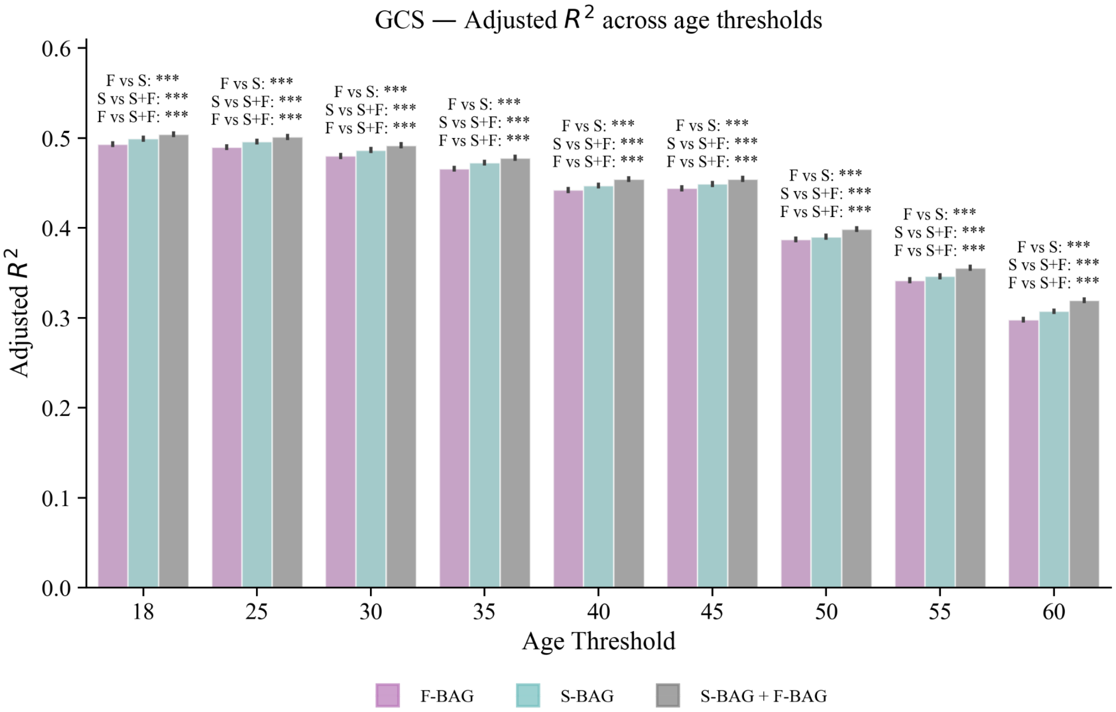

**Figure S5.1: GCS regression performance across age thresholds.** GCS regression performance is shown for the following models: F-BAG (F), S-BAG (S) and S-BAG + F-BAG (S+F). All models account for age, age-squared, sex and education as covariates. Error bars represent the 95% CI across 1000 subsamples (n=400 subjects). All statistical tests significance are annotated for each age threshold. ***: $p<0.001$

##### 2. Covariates role in GCS regression across age thresholds

OLS regression of GCS has been done from S-BAG and F-BAG combined age, age-squared, sex and education as covariates. Fig. S5.2 show the mean coefficient across 1000 subsamples (n=400 subjects per subsample), annotating "*" if the effect was found to be robust (see Methods for details). Education is the only covariate being always positively contributing to the prediction, meaning that higher education was associated with higher global cognitive score. Age-squared had a robust effect only when younger subjects were included, as expected from the GCS decline with age. Age was only robust for the age threshold of 45 years. Sex had a robust effect later in life, with very low contribution to the prediction.

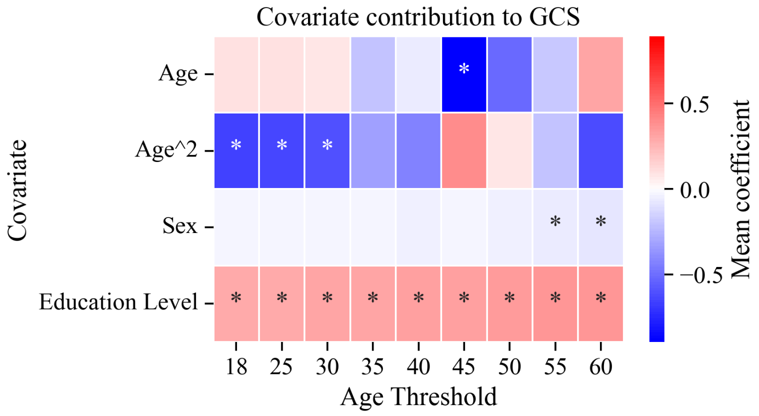

**Figure S5.2: Mean coefficient for each covariate across age thresholds.** * is present if the effect is considered to be robust across 1000 subsamples (n=400 subjects per subsample).

**3. GCS regression from original brain features**

Here we predicted GCS directly from regional brain features using Ridge regression. All models included chronological age, age-squared, sex, and education as covariates and were evaluated using stratified 5-fold cross-validation. Separate models were fitted for cortical thickness, volume, SC strength, and FC strength. Each model included 100 regional features plus the four covariates. In the full sample, cortical thickness predicted GCS with a cross-validated $R^2$ of 0.461 ± 0.063, volume with $R^2$ = 0.467 ± 0.036, SC strength with $R^2$ = 0.429 ± 0.055, and FC strength with $R^2$ = 0.467 ± 0.036. In Fig. S5.3 the regional coefficients for each modalities are shown along the S-A axis, on the Schaefer parcellation and with the average values per resting-state network.

We additionally fitted a combined connectivity-strength model including both SC strength and FC strength, for a total of 200 regional features plus covariates. This model reached a cross-validated $R^2$ of 0.411 ± 0.041 in the full sample. Performance decreased across age-restricted samples, with $R^2$ = 0.391 ± 0.029 for participants older than 45 years, $R^2$ = 0.311 ± 0.032 for participants older than 50 years, $R^2$ = 0.265 ± 0.071 for participants older than 55 years, and $R^2$ = 0.194 ± 0.114 for participants older than 60 years. To compare structural and functional connectivity, we directly compared SC strength and FC strength models. Both models included 100 features, the same covariates, the same participants, and the same cross-validation scheme. FC strength consistently outperformed SC strength across all age thresholds. Specifically, FC strength reached $R^2$ = 0.467, 0.428, 0.379, 0.314, and 0.274 across the full, >45, >50, >55, and >60 samples, respectively. In contrast, SC strength reached $R^2$ = 0.429, 0.394, 0.325, 0.250, and 0.169 across the same age thresholds.

Overall, regional brain features contained cognition-relevant information, with FC strength showing the strongest performance among the connectivity-strength models. However, these high-dimensional regional models did not outperform the BAG-based OLS model, suggesting that S-BAG and F-BAG provide a compact and interpretable summary of cognition-relevant brain variability.

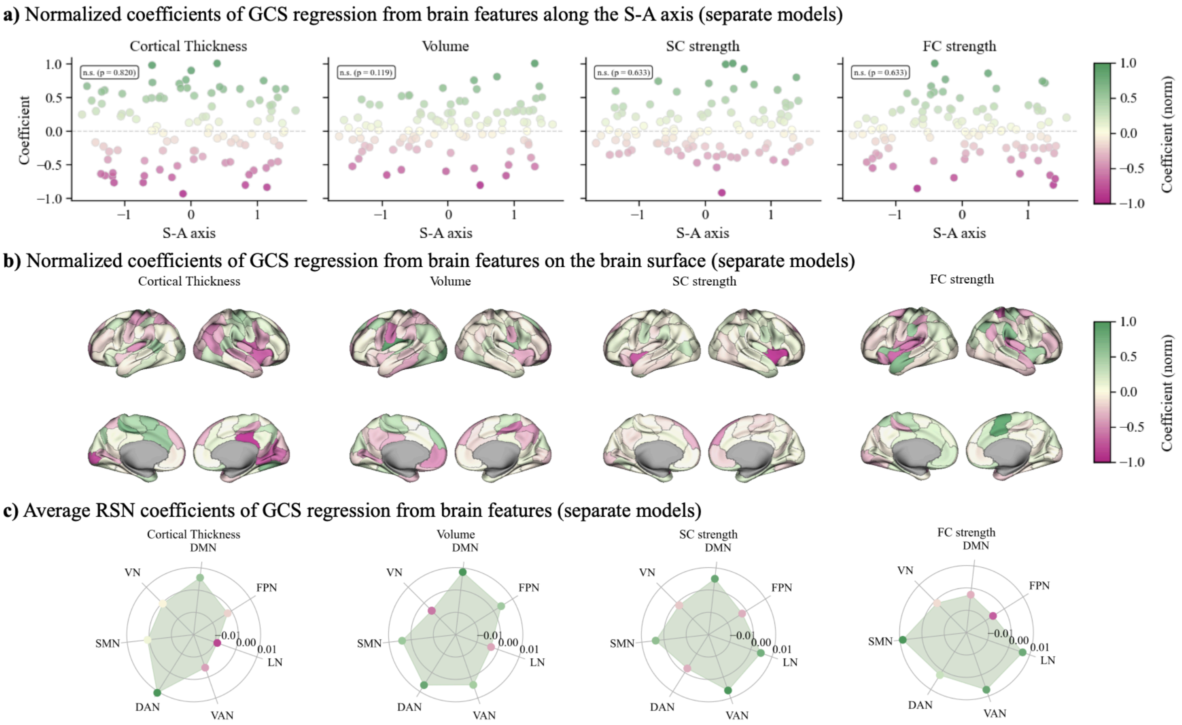

**Figure S5.3: GCS regression from different modalities. a)** ROI-based regression coefficients along the sensorimotor-association (S-A) axis. No significant association with the S-A axis has been found. **b)** ROI-based regression coefficients on the 100 ROIs Schaefer atlas. Positive values (greener) indicate positive coefficients. Negative values (pinker) indicate negative coefficients. c) Average coefficients per resting-state network.

**4. Additive models were preferred over interaction models**

Additive and interaction models were compared using the Bayesian Information Criterion, with lower BIC values indicating better model fit after penalizing model complexity. All models were fitted in the subsample of participants older than 45 years, in line with the age range in which age-related cognitive variability is expected to become more pronounced. Model coefficients, p-values, $R^2$, and BIC values were reported for each cognitive outcome.

We also tested whether cognition was better explained by an interaction between structural and functional brain aging by adding an S-BAG × F-BAG interaction term to each domain-specific model. Adding the S-BAG × F-BAG interaction term did not improve model fit: across the global cognitive score and all cognitive-domain targets, the additive model consistently showed lower Bayes Information Criterion (BIC) values than the interaction model. The interaction coefficient was not significant for five out of six outcomes. This suggests that the interaction effect was small and did not justify the inclusion of an additional parameter. This indicates that S-BAG and F-BAG contributed to cognition primarily in an additive, rather than interactive, manner. Findings support the use of additive models and suggest that structural and functional brain age gaps contribute to cognitive variability primarily through partially independent rather than multiplicative effects.

#### 6. Parcellation robustness

To better assess the reliability of the results on the 100 Schaefer atlas, we also considered the 400 ROIs Schaefer parcellation. We first constructed the brain age model from structural (cortical thickness, volume and SC strength combined) and functional (FC strength) features using the Ridge regression. In both cases the 400 ROIs atlas achieved a slightly higher accuracy with respect to 100 ROIs, but with higher overfitting on the train set, especially for the functional age prediction (Fig. S18a).
Then, after correcting for the age-bias of the predictions, we saw that S-BAG and F-BAG were similarly associated in the two parcellations, with slightly higher association for 4000 ROIs, probably due to the higher accuracy (Fig. S6b). Moreover, both BAG markers show reproducible correlation with the GCS (Fig. S6c,d). Note that while F-BAG has lower association in the 400 ROIs, supporting the idea that increasing accuracy does not always reflect in higher sensitivity of BAGs.
Finally, we considered the BAG markers to predict the GCS across age thresholds (Fig. S6e), and to predict the cognitive domains (Fig. S6f), accounting for age, age-squared, education and sex. All cases show comparable R-squared between parellations. Overall, the S-BAG showed similar or greater contribution in the 400 ROIs atlas (Fig. S6g), while the F-BAG was weaker with respect to the 100 ROIs atlas (Fig. S6h). This effect is in line with BAG markers correlation with the GCS. Moreover, since the 400 ROIs brain age models showed greater overfitting, results could be less reliable for the more refined parcellation. This finding is in line with All covariates showed comparable contribution across the two atlases (Fig. S6i-l).

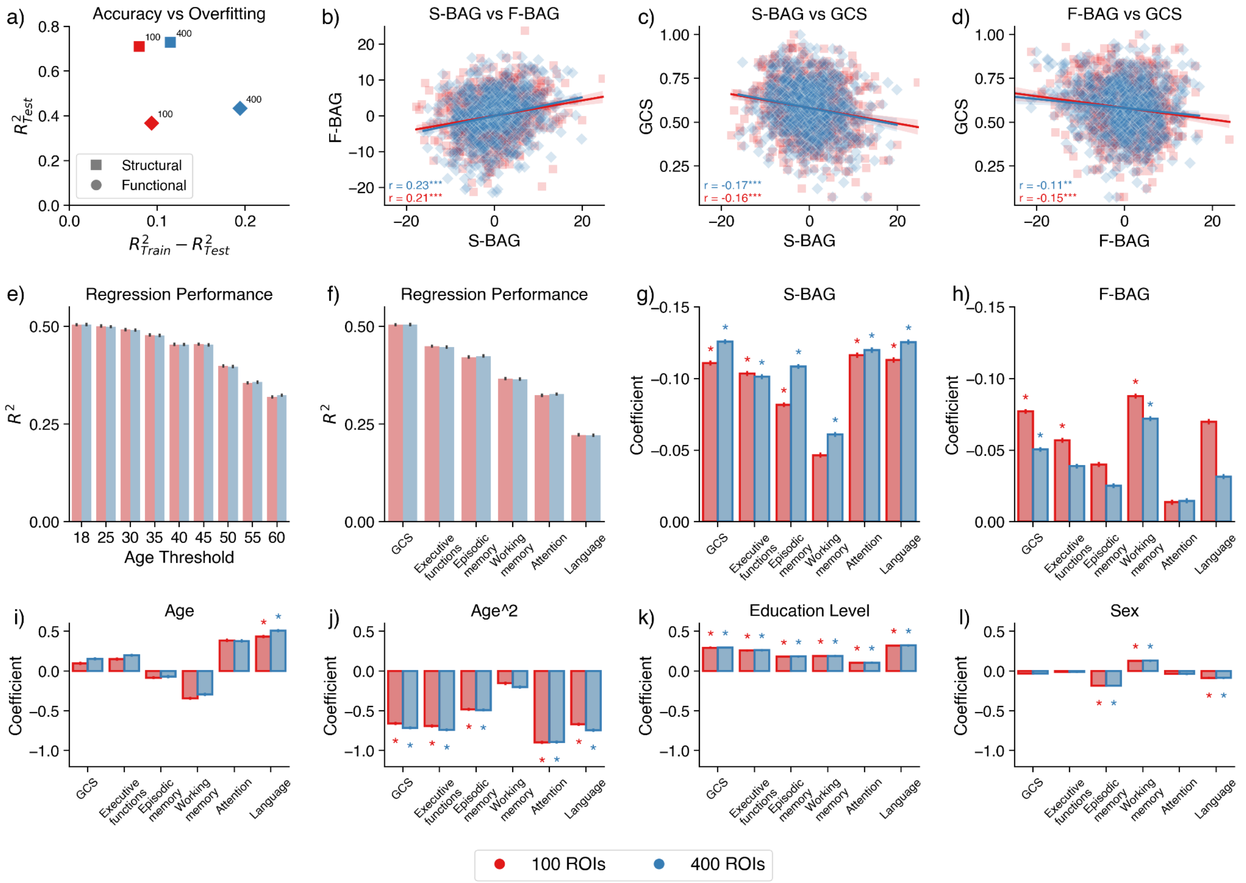

**Figure S6: Results comparison between 100 (red) and 400 (blue) ROIs Schaefer atlases. a)** Brain age prediction accuracy (R-squared of test set) as a function of the overfitting (difference between train set and test set R-squared) for both structural and functional models. b) Correlation between S-BAG and F-BAG with respective Spearman's correlation. r=0.21 ($N_{ROIs}$=100), r=0.23 ($N_{ROIs}$=400). ***: p<0.001, **:p<0.01. c) Correlation between S-BAG and GCS with respective Spearman's correlation. r=-0.16 ($N_{ROIs}$=100), r=-0.17 ($N_{ROIs}$=400). ***: p<0.001, **:p<0.01. d) Correlation between F-BAG and GCS with respective Spearman's correlation. r=-0.15 ($N_{ROIs}$=100), r= -0.11 ($N_{ROIs}$=400). ***: p<0.001, **:p<0.01. e) GCS regression performance across age thresholds. f) Domain regression performance on full sample. g) S-BAG contribution to domain regression. h) F-BAG contribution to domain regression. i) Age contribution to domain regression. j) Age-squared contribution to domain regression. k) Education contribution to domain regression. l) Sex contribution to domain regression. * is present if the contribution effect is considered robust across 1000 subsamples (n = 400).
